# Robust and scalable single-molecule protein sequencing with fluorosequencing

**DOI:** 10.1101/2023.09.15.558007

**Authors:** James H. Mapes, Julia Stover, Heather D. Stout, Tucker M. Folsom, Emily Babcock, Sandra Loudwig, Christopher Martin, Mariah J. Austin, Fan Tu, Casey J. Howdieshell, Zachary B. Simpson, Thomas Blom, Daniel Weaver, Daniel Winkler, Kent Vander Velden, Parham M. Ossareh, John M Beierle, Talli Somekh, Angela M. Bardo, Eric V. Anslyn, Edward M. Marcotte, Jagannath Swaminathan

## Abstract

The need to accurately survey proteins and their modifications with ever higher sensitivities, particularly in clinical settings with limited samples, is spurring development of new single molecule proteomics technologies. Fluorosequencing is one such highly parallelized single molecule peptide sequencing platform, based on determining the sequence positions of select amino acid types within peptides to enable their identification and quantification from a reference database. Here, we describe substantial improvements to fluorosequencing, including identifying fluorophores compatible with the sequencing chemistry, mitigating dye-dye interactions through the use of extended polyproline linkers, and developing an end-to-end workflow for sample preparation and sequencing. We demonstrate by fluorosequencing peptides in mixtures and identifying a target neoantigen from a database of decoy MHC peptides, highlighting the potential of the technology for high sensitivity clinical applications.

## INTRODUCTION

Modern diagnostics have made significant progress by determining the abundance of individual proteins, as exemplified by detecting prostate-specific antigen with antibody tests (Thompson et al. 2004) or quantifying PD-L1 levels to stratify patients for immunotherapy (Kowanetz et al. 2018). However, many diseases arise due to perturbations in the levels of multiple proteins, their modifications and their interactions, making it crucial to accurately measure them simultaneously for diagnosis or treatment. Historically, proteomics technologies have required either minimal sample abundances of proteins in the range of femto-to attomole, e.g. in the case of typical mass spectrometry experiments (Timp and Timp 2020), or suffered from low quantitative accuracy, as in the case of affinity-based methods (Katz et al. 2022). Applications to clinical settings in particular could benefit from technology capable of accurately quantifying multiple proteins in their various modified states with high sensitivity, while working with limited samples, such as small tissue biopsies or fine needle aspirates (Boys et al. 2023).

This need for quantitative, high-sensitivity measurements of peptides and proteins from biological mixtures has recently spurred the development of new single molecule protein sequencing technologies (Restrepo-Pérez, Joo, and Dekker 2018; Floyd and Marcotte 2022; Brady and Meyer 2022; Callahan et al. 2020; Alfaro et al. 2021). These technologies are currently in various stages of development, from conceptual proposals (Palmblad 2021; Egertson et al. 2021) to demonstrations on simple peptide or protein samples (Reed et al. 2022; Brinkerhoff et al. 2021; Aubin-Tam et al. 2011; de Lannoy et al. 2021; Borgo and Havranek 2014), but in general have not yet been applied to analyses of complex peptide mixtures. While the current inability of these technologies to scale to complex samples can be ascribed to the natural pace of early technological advancements, a number of challenges still need to be overcome in order to achieve general utility.

We previously demonstrated a highly parallelized single molecule peptide sequencing platform called *fluorosequencing* (Swaminathan et al. 2019). Fluorosequencing is based on the principle that determining the positions of a few, select amino acids within a peptide can be sufficient for matching the partial sequence (termed a *fluorosequence*) to a reference protein set to infer the identity of the peptide or protein. To determine the fluorosequence for individual peptide molecules, first, peptide side chains of select amino acid types are labeled with fluorescent dyes. Millions of these labeled peptides are then immobilized in a flow cell and imaged using total internal reflection fluorescence (TIRF) microscopy. Cycles of Edman degradation are performed, which removes one N-terminal amino acid from each peptide on each cycle, and the peptides’ fluorescent intensities measured after each Edman cycle. Using image analysis and signal processing, the cycles corresponding to the removal of fluorescent amino acids are determined for each molecule, resulting in a fluorosequence for each molecule that can be matched to the reference database to identify the peptide. In prior publications (Swaminathan, Boulgakov, and Marcotte 2015; Swaminathan et al. 2019), we introduced this concept and methodology, discussed the sources of errors, and demonstrated its feasibility for sequencing simple peptide mixtures. However, our data was mostly limited to a single fluorescent channel.

Here, we describe significant improvements to fluorosequencing and detail the challenge of mitigating dye-dye interactions encountered during scaling of the technology. We introduce our solution to minimize these interactions through the creation of rigid polyproline fluorophore linkers, and we experimentally demonstrate a robust and scalable process for identifying peptides in a complex mixture. Finally, we showcase the targeted ability of the fluorosequencing technology to identify a putative antigenic HLA-1 peptide in a reference background of 37 alternative HLA-1 peptides, suggesting the utility of this technology in clinical applications.

## RESULTS

### Improvements to the end-to-end process

We made significant improvements to the technology in three major areas: (a) the *sample preparation*, specifically the chemistry of labeling peptides, (b) peptide *sequencing*, consisting of alternating cycles of Edman degradation and single molecule imaging, and (c) the *computational analysis*, especially the image processing and peptide-read mapping. A glossary of terms and the peptides analyzed (named with the convention *JSP#*) are provided in the **Supplementary Tables ST1 and ST2). Fig. 1** summarizes the changes and improvements to the process, which we next describe in detail.

**Fig. 1:**
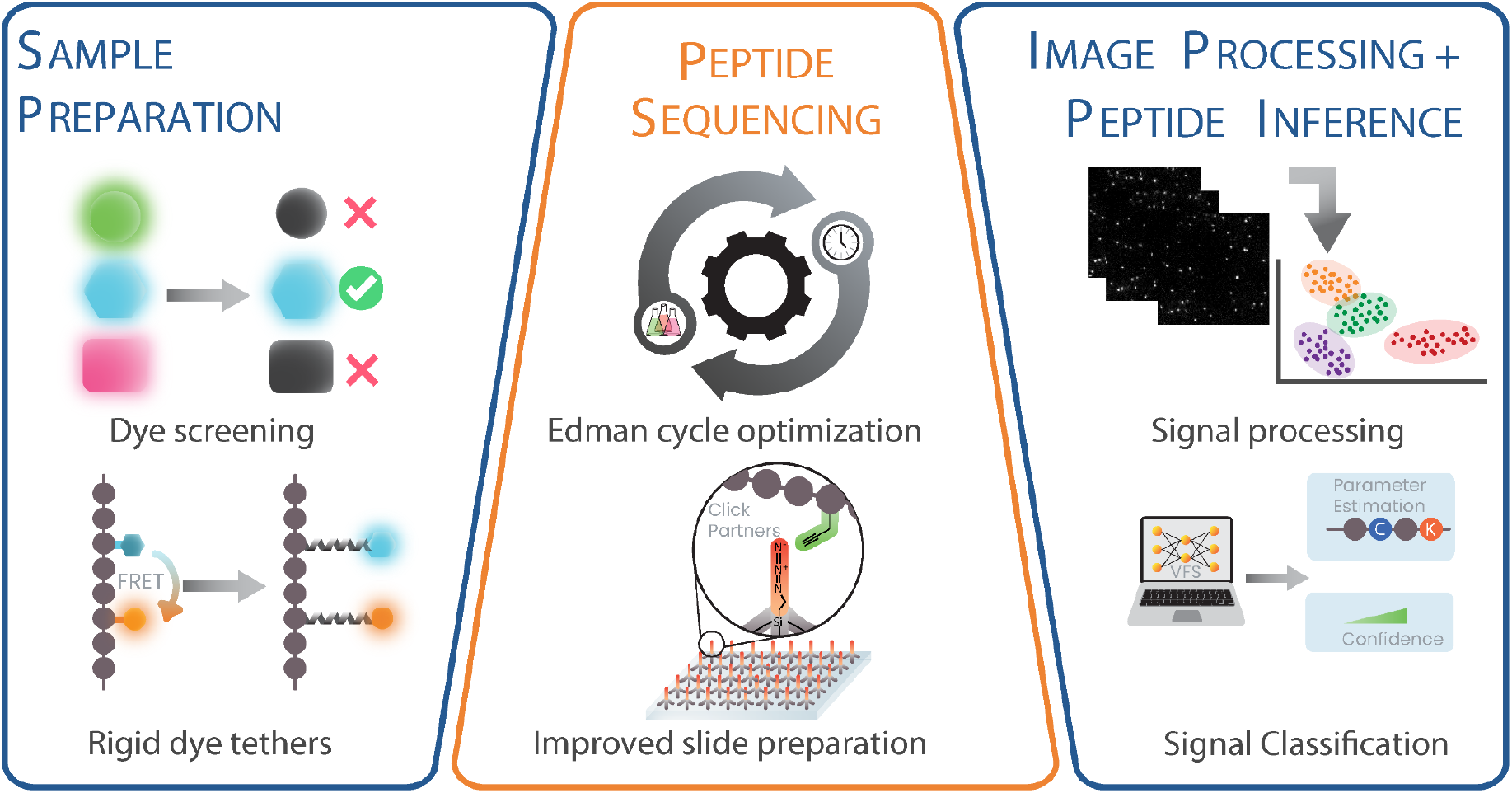
Overview of fluorosequencing technology highlighting the improvements and developments to the process. We substantially improved the sample preparation, imaging, fluidics, image processing, and peptide-read matching workflows. To improve sample preparation, we screened 70 dyes for improved photophysical properties and chemical stability to Edman solvents, selecting dyes across 6 fluorescent channels. We mitigated dye-dye interactions on fluorescently labeled peptides through spacing fluorophores from the peptide backbone using long polyproline rigid tethers. We optimized solvents and conditions for increasing Edman efficiency in less time, and also decreased non-specific binding to flow cell surfaces by 23-fold by immobilizing peptides on azide-derivatized surfaces using click chemistry. Finally, we developed and implemented a scalable image processing pipeline and a novel machine learning classifier framework to infer peptide identities from raw reads.

### Expanding the set of fluorescent dyes compatible with fluorosequencing

Based on computational simulations of fluorosequencing (Swaminathan, Boulgakov, and Marcotte 2015), the identification of proteins in complex samples will generally require selectively labeling 3 or 4 amino acid types, each with a different fluorophores. We previously observed that commonly used fluorophores, such as BODIPY and cyanine dyes, did not recover fluorescence after exposure to the solvents and reagents used in Edman sequencing chemistry, limiting the number of distinct fluorescent labels available for sequencing (Swaminathan et al. 2019). Therefore, to identify additional fluorophores compatible with Edman sequencing chemistry, we conducted a screen of 70 fluorescent dyes, spanning both commercially-available and custom-synthesized dyes, and found 31 with the required stability in critical Edman reagents. For this screen, we immobilized the succinimidyl variants of dyes on amine-functionalized Tentagel beads and measured fluorescence recovery after exposure to methanol, trifluoroacetic acid, or 20% phenylisothiocyanate in pyridine. We also tested for stability in piperidine solution to ensure compatibility with synthetic peptides, which often require removal of an N-terminal Fmoc blocking group prior to fluorosequencing. **Supplementary Table ST3** provides the solvent stability data for each of the fluorophores we tested. We observed that rhodamine and carbopyronine dyes generally showed superior stability to the Edman reagents.

From this set of s-stable dyes, we further prioritized dyes based on their practical performance in fluorosequencing experiments. We measured four different experimental parameters for the dyes, including their *chemical destruction rate*, the per cycle fluorescence loss of dyes on N-terminally acetylated (hence, not sequenceable) peptides after exposure to Edman reagents; *photobleaching rate*, the fluorescence loss rate during constant illumination in imaging solvent (0.1 mM Trolox in degassed methanol); and their fluorescence *brightness* and its *variation*. **Supplementary Fig. 1A** reports these parameters for two of the dyes we selected, TexasRed and Atto643. We measured photobleaching and chemical destruction rates for Atto643 at 1.9% and 2% per cycle, respectively, and TexasRed at 0.46% and 5.6% per cycle. For the rest of this work, we generally use TexasRed, Alexa555, and Atto643 as a reasonable starting palette of fluorosequencing labels. **Supplementary Table ST4** reports their relevant parameters, while **Supplementary Fig. SF1B** highlights four additional dyes that also appeared suitable for fluorosequencing experiments.

### Alkyne modifications to peptides reduce non-specific peptide binding to the silane surface

A second major improvement is in how peptides are anchored in the flow cell for sequencing. We previously observed significant amounts of non-specific binding of labeled peptides to aminosilane-labeled glass surfaces, an effect we attempted to control for by fluorosequencing negative control (N-terminally acetylated) peptides in parallel (Swaminathan et al. 2019). We speculated that non-specific interactions of either the peptide or the dye with the slide surface might be due to the charged amine groups of the 3-Aminopropyl)triethoxysilane, and that by moving to an uncharged surface, such as by immobilizing peptides using azide-alkyne click chemistry, we might reduce the observed nonspecific binding (Charlton et al. 2011). We therefore functionalized glass slides with 3-azidopropyltriethoxysilane and modified peptides to contain C-terminal alkyl groups. For synthetic peptides, this was achieved using Fmoc-propargylglycine as the C-terminal amino acid building block. After making this modification we observed a 23-fold improvement in correctly sequenceable peptides (Hinson et al. 2021). We therefore moved exclusively to immobilizing all peptides in this work by azide-alkyne click chemistry.

### Changes to Edman chemistry increase the efficiency and speed and reduce dye destruction rates

Next, we sought to make improvements to the slide preparation process and the Edman sequencing chemistry itself, and thus further reduce the rates of slide surface and dye destruction due to the sequencing chemistry (E. A. Smith and Chen 2008). First we explored the role of incubation times of the various Edman reagents. In particular, changing the TFA incubation times caused a dramatic improvement to our previously reported sequencing efficiency, as shown in **Supplementary Fig. SF2A**. We choose to set the TFA incubation time to 8 mins per cycle, as longer incubation times caused unacceptable loss of dye fluorescence

Similarly, we found that doubling either the incubation time or concentration of the Edman reagent, phenylisothiocyanate, improved our efficiency; moving forward, we used 20% v/v in pyridine (twice our prior concentration) to avoid increasing reaction times. Additionally, we found that adding 60mM of N-methylmorpholine base into the phenylisothiocyanate coupling solution and increasing the reaction temperature by 10°C further increased our reaction efficiency (**Supplementary Fig. SF2B**). Applying these changes improved the portion of peptides showing the expected fluorosequence (an easily-measured metric that correlates with Edman efficiency) for a difficult to sequence proline-containing peptide (*JSP263*) from 35 to 67% (**Fig. 2A**), corresponding to ∼91-99% Edman efficiency. We saw similar improvements across a wide variety of peptides of varying composition, as shown for several examples in **Fig. 2B**. In general, we found proline to be the most resistant to Edman cleavage, consistent with historic observations (Brandt et al. 1976), with an average decrease of 9.5% in the percent of peptides successfully showing the largest fluorescent intensity decrease at the expected label position when the labeled residue is preceded by proline (**Supplementary Fig. SF3**). The effect depended in part on the identity of the amino acid that follows the proline, likely due to the high energy barrier in forming the bicyclic of phenylthiohydantoin-proline during the Edman degradation mechanism (Tarr 1975; Brandt et al. 1976). As a final improvement to the Edman sequencing process, we simplified the fluidic system by eliminating the free-base step and substituting ethyl acetate with acetonitrile. We also observed that water in the base mix is critical for successful Edman chemistry and purging with nitrogen has only limited benefits. With all of these changes considered together, the full cycle of Edman chemistry has now been reduced from 75 minutes to 40 minutes and an average increase in Edman efficiency to 95-99%.

**Fig. 2:**
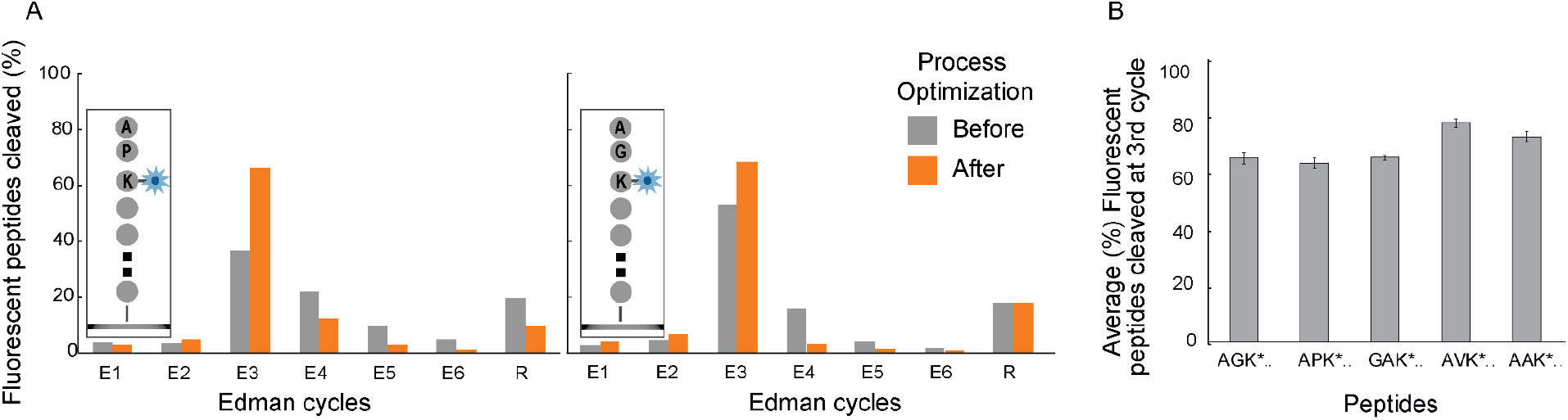
Optimization of Edman conditions and solvents increased the Edman efficiency across multiple amino acids and peptide sequences. (A) Lysines at the third amino acid sequence position were fluorescently labeled in two peptides that contained either a preceding N-terminal proline (in JSP263; left panel) or glycine (in JSP254; right panel) residue. Following Edman optimization procedures, the largest fluorescence intensity drop per molecule occurred at the expected sequence positions in 63% and 74% of peptides, respectively. (B) We observed an average of 63-75% of peptides with differing N-terminal amino acid sequences correctly showing the largest fluorescence intensity drop at the labeled lysine residue. These rates correspond approximately to an Edman efficiency of 91-99% per cycle.

### Vapor deposition of silane improves consistency and reduces fluorescent contamination on flow cell surfaces

We also sought to improve the quality of surface preparations in the sequencing flow cells, as we often noted contaminating fluorescent molecules in early sequencing experiments. We therefore implemented a vapor deposition method for applying silane to the glass flow cell surface instead of the previously used dip coating method. This change resulted in a significant reduction of fluorescent contaminants in the 488, 532 and 561 nm imaging channels, which we ascribe to reduced handling, and increased uniformity of the surfaces and across preparation batches due to reduced silane polymerization (Popat, Johnson, and Desai 2002). **Supplementary Fig. SF4** illustrates the in-house vapor deposition method and highlights the quality of the resulting surfaces.

### Dye-dye interactions present challenges to identifying label positions

With all these optimizations in place, sequencing quality was significantly improved. However, as we began installing multiple numbers and types of fluorophores on peptides, we increasingly observed dye-dye interactions that complicated deconstructing the observed fluorescent signals into their respective counts and types of fluorophores, a critical interpretive step. We observed two different types of dye-dye interactions in our system - dye quenching and Förster resonance energy transfer (FRET).

Dye quenching was evident when multiple copies of the same Atto647N fluorescent dye were installed on the same peptide. For example, we observed that the fluorescence intensity distribution of peptides containing either 1, 2, 3, or 4 Atto647N dyes did not follow a simple additive increase of single dye intensity distribution (**Supplementary Fig. SF6)**. This behavior could be attributed to dye quenching between closely spaced dyes.

In contrast, FRET was evident when imaging peptides with two different fluorophores, in which we observed signals primarily from the higher wavelength emitter. Prior to optimizing the sequencing chemistries and surfaces, earlier sequencing experiments showed relatively little evidence for such interactions; we speculate that high dud-dye rates (see glossary in **Supplementary Table ST1)** and non-specific binding of dyes to the surface conflated the signal. Moreover, we suspected that FRET might also be unlikely due to the flexibility of the fluorophore linker attached to the peptide backbone and the use of organic solvents (methanol) for imaging, which prohibits dye-dye interactions such as pi-stacking and H-dimer formation (Ogawa et al. 2009). However, with the system-wide improvements made to reduce background contamination, an effect due to FRET was considerably more evident.

As an example of the FRET observed, **Supplementary Fig SF7A** shows data for a peptide (JSP129) containing two distinct fluorophores, Janelia Fluor 549 and Atto647N, with less than 5% colocalization of peptide peaks from the two imaging channels. To explore if this observation resulted from FRET, we imaged fluorescent single molecules using the expected excitation and emission for each fluorophore and an additional “FRET channel”, exciting with the lower wavelength and measuring the higher wavelength emission. For a majority of the peptides, we observed significant signals in the FRET channel and the Atto647N channel; bleaching the Atto647N caused the Janelia Fluor 549 signal to return, confirming intramolecular dye-dye energy transfer. We concluded that FRET occurred at a very high rate in this and other peptides, with a measured FRET efficiency of >90% for this pair of dyes (see Methods). We observed similarly low colocalization and high FRET between the dye pairs Alexa488/Atto647N and JF525/Atto647N (**Supplementary Fig. SF7B**). Thus, we made a major effort to try to mitigate both types of dye-dye interactions by introducing the appropriate linkers for attaching the dyes to the peptides, intended to position the dyes far enough apart to reduce their interactions.

### Rigid polyproline linkers (*Promers*) minimize dye-dye interactions

Previously, we found that attaching fluorophores to peptides using flexible (PEG)_10_ linkers provided a modest reduction of dye-dye quenching (Bachman et al. 2022). However, we suspected that the short length (approx. 30 Å) and high flexibility of these PEG polymers limited their effectiveness in spatially separating the fluorophores. To test this hypothesis, we synthesized a branched peptide with two 14 subunit polyproline peptides on the two arms, each labeled with either TMR or Atto647N. Our observation of a 39% recovery of the donor signal in this system (see **Supplementary Fig. SF8A**), when compared to a peptide system without the polyproline linker, indicated that spacing dyes through rigid polyproline linkers might be a viable approach to mitigate dye-dye interactions.

To explore this notion more fully, we tested several different length proline linkers and determined that a 30-proline linker significantly reduced FRET to less than 10% while still being synthetically accessible (**Supplementary Fig. SF8B**). To install these long polyproline linkers with fluorophores on peptides, we designed and synthesized peptidic linkers (termed “Promers”) with the following characteristics: At the N-terminus, we synthesized a “Gly-(PEG)_2_-Gly” unit to increase solubility and flexibility, followed by a 30-proline repeat. The terminal amine was labeled with a reactive chemical moiety, such as dibenzocyclooctyne (DBCO), and at the C-terminus, we synthesized with a lysine, whose side chain could then be functionalized with the desired fluorophore **(Fig. 3A**). In general, polyproline is thought to organize as a rigid rod (Schuler et al. 2005; El-Baba et al. 2019) whose length, for the case of the Promers and depending on the polarity of the solvent, is estimated to be between 6-9 nm, above the Förster radii of many fluorophore pairs.

**Fig. 3:**
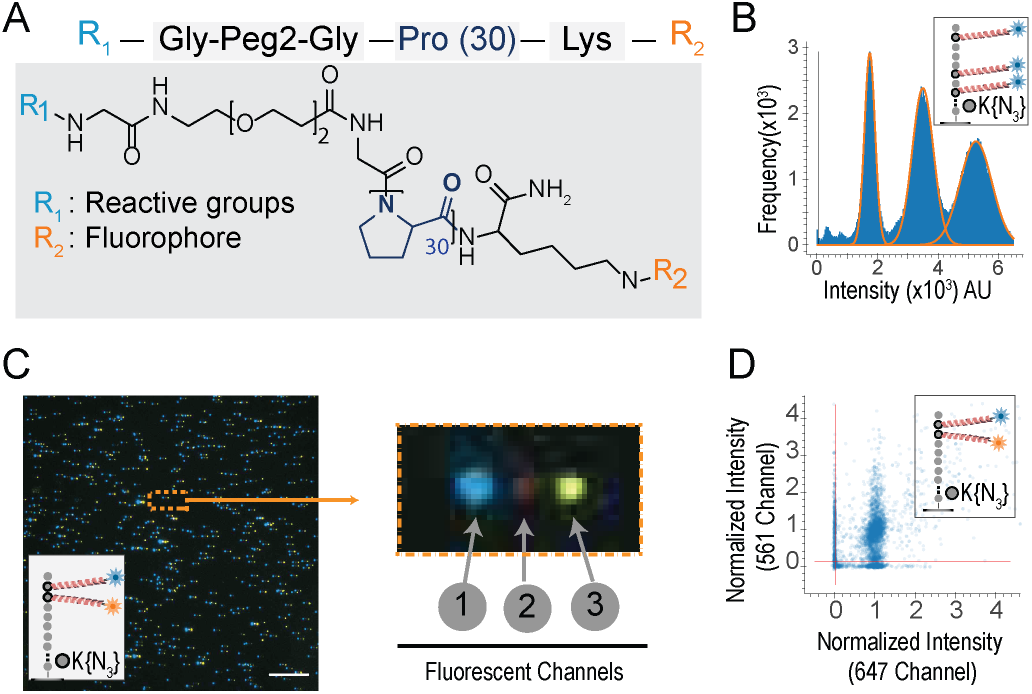
PEG/polyproline linkers (Promers) mitigate dye-dye interactions on peptides with multiple fluorophores. (A) illustrates Promer design and structure, with a 30-unit proline repeat flanked by a fluorophore (R2) linked to a lysine residue, and a DBCO reactive moiety (R1) linked *via* a flexible Glycine-PEG2-Glycine spacer. (B) The intensity histogram of 59,405 partially photobleached peptides (*JSP126*) shows three distinct peaks, indicating the resolution of one, two, or three active Atto643 fluorophores (installed *via* Promers at azido-lysine residues at amino acid positions 2, 6, and 8). The additive nature of the fluorescence intensities (median values for 1, 2, and 3 dyes, respectively, are 17,838; 35,040; and 52,242 arbitrary units) demonstrates minimal quenching between the fluorophores. (C) A representative TIRF micrograph (left panel) of individual *JSP212* peptides, labeled with Atto643 (FRET acceptor) and JF549 (FRET donor) dyes on Promers at the 2nd and 3rd amino acid positions, demonstrates low FRET levels between the dyes. This composite image was made from offsetting the signal from three fluorescent channels (1-acceptor, 2-FRET, and 3-donor), shown enlarged for a single peptide molecule at right. The presence of signals in both donor and acceptor channels indicates the magnitude of FRET effect is less than 10% even on consecutive amino acids due to the Promers. (D) A scatter plot of fluorescence intensities across the donor and acceptor channels shows that 67% of individual peptide molecules (10,385 filtered peptides) exhibit two distinct fluorophores, confirming low FRET levels when dyes are tethered by Promers. Scale bar, 10 μm.

These Promers allowed us to investigate the influence of quenching and FRET on peptides with multiple fluorophores. To examine the occurrence of quenching, we installed three Atto643 labeled Promers on the peptide (*JSP216*), and through multiple fields of imaging, could clearly discriminate the intensity of peptides with one, two and three fluorophore signals (**Fig. 3B**). The peaks’ mean intensities of 17,472; 34,521; and 52,381 arbitrary fluorescence units closely match an additive model of a single fluorophore’s intensity. Next, to evaluate a potential reduction in FRET, we synthesized a peptide (*JSP212*) containing two distinct fluorophores, Janelia Fluor JF549 and Atto643, attached *via* Promers. We observed less than 10% FRET efficiency and 67% colocalization of spots for two fluorophore peptide (*JSP212*) between the donor, JF549 and acceptor Atto643, channels (**Fig. 3C**). Labeling peptides with Promers did not impact the Edman efficiency (**Supplementary Fig. SF9**). Thus, by substantially mitigating dye-dye interactions, Promers help recover signal lost to FRET or quenching and consequently improve fluorosequencing data quality, as we discuss next.

### Identifying peptides in a mixture using an end-to-end workflow

As a major goal of these improvements is to expand fluorosequencing to increasingly diverse samples and mixtures, we tested the improved workflow and labeling chemistry for its performance at identifying peptides in a mixture. We began by fully characterizing the fluorosequencing of four individual synthetic peptides with similar sequences, with each peptide having 2 or 3 amino acids labeled with either Atto647N or Alexa555 Promers (**Supplementary Table ST5**). As a guide to their subsequent interpretation, we also determined key experimental parameters from these and other control experiments, including Edman efficiency, dud-dye rates, and peptide detachment rates, in addition to those rates in **Supplementary Table ST4**. These rates are a key element of computational models for interpreting the fluorosequencing datasets, as we discuss below.

We then considered the peptides as a mixture: we combined the four synthetic peptides in an approximate equimolar ratio and performed 15 rounds of fluorosequencing, collecting 100 fields of images across the two fluorescent channels (**Fig. 4A**). The left panel of **Fig. 4B** shows a representative image of the 4 peptide mixture, overlaid with two fluorescent channels and indicates positions of four distinct peptide molecules (labeled 1-4). The right panel shows expanded images of individual fluorescent peaks for the four peptides across the two fluorescent channels after each cycle of Edman chemistry, where stepwise reductions in fluorescent intensity can be readily observed. The images were then processed and the reads extracted and filtered (see **Methods**) using a computational workflow (illustrated in **Supplementary Fig. SF5)**. This signal processing workflow involves analyzing TIRF microscope images to identify peaks and calculate each peak’s intensity at the end of every cycle, thus generating an intensity track from all fluorescent channels (*raw sequencing reads)*.

**Fig. 4:**
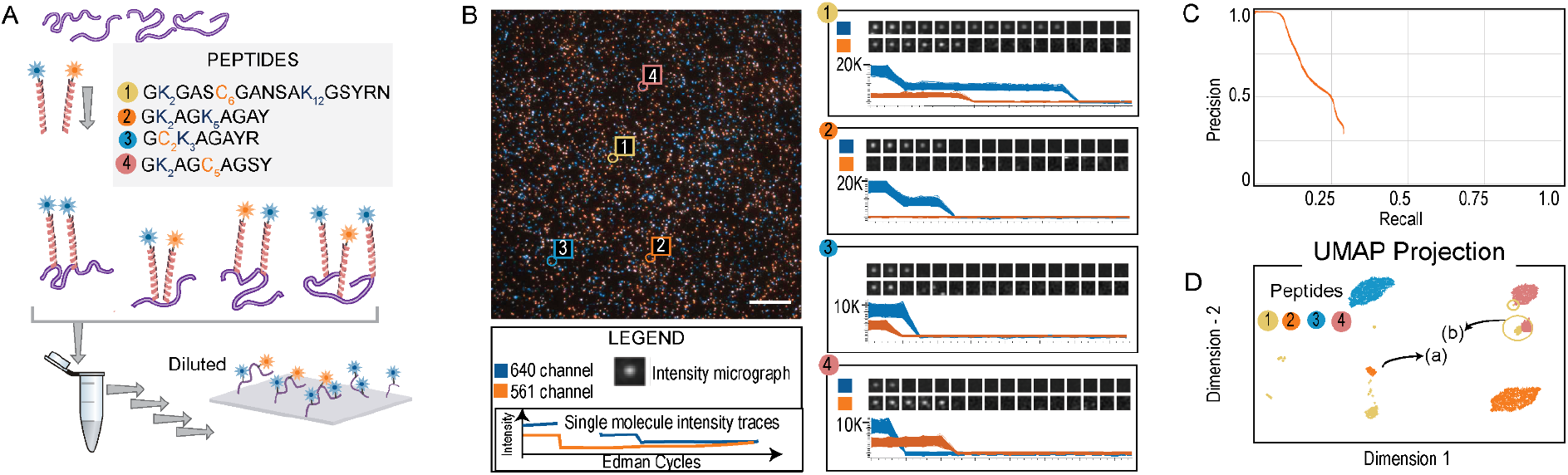
Fluorosequencing a two color, four peptide mixture. (A) Four fluorescently labeled peptides (detailed in **Supplementary Table ST5**) were mixed at approximately equimolar (2 μM) concentrations, diluted by four orders of magnitude (to 200 pM), and fluorosequenced, collecting raw sequencing reads for 49,480 individual peptide molecules. (B) shows a representative TIRF microscope image with overlaid 561 and 647 nm channels, with data plotted at right for four distinct peptides labeled 1-4, each exhibiting a unique fluorosequencing profile, shown in the individual peptide micrographs for consecutive Edman cycles in the two fluorescent channels. The associated plots report sequencing read fluorescent intensities for 331, 1423, 1731, and 1012 replicate molecules, respectively. (C) Using a machine learning classifier, peptides can be accurately classified to one of the 4 source peptides in a reference database with 50 decoy peptides. The 4,948 peptides classified with scores >0.99 were also visualized in a UMAP plot in (D) by considering their raw reads as feature vectors; the four distinct peptides are clearly separable. Also shown is the presence of peptide-2 with a missed Edman cycle, depicted in cluster (a), but which is still classified correctly. A minor population of peptides with photobleached Atto647 dye, depicted in cluster (b), was classified as peptide-1.

Peptide inference (in particular, *peptide-read matching*) can then be performed using a machine learning classifier, trained to assign reads to peptides from the reference database based on simulating fluorosequencing with the same experimental parameters (efficiency of Edman cleavage, photobleaching rate, dud-dye rates, chemical destruction rates, etc). This assigns each read to a reference database peptide with an associated confidence score. A detailed description along with pseudocode is provided in **Supplementary Information**. For the case of the four peptide mixture, we considered a reference peptide database of 54 peptides - containing the four synthetic peptides to be sequenced and an additional 50 decoy peptides, a randomly chosen set of 20-amino-acid long peptides. For computational modeling and inference purposes, we changed the azido-lysine on the input peptide sequence to cysteine. These simulated data were used as the training set for a random forest classifier (see Methods) for identifying the real peptides from the larger reference database. We also subsequently develop more scalable approaches to solve this problem (M. B. Smith, Simpson, and Marcotte 2023).

Using the machine learning classifier, we scored each fluorosequencing read to the most likely source peptide in the reference database. As plotted in **Supplementary Fig. SF14A**, scores above 0.99 predominantly identified the true peptides. Plotting the corresponding Precision-Recall curve indicates that 6% of the raw reads can be correctly classified with 99% precision to the four input peptides (**Fig. 4C)**. Visualization of the high scoring reads in a UMAP plot (McInnes, Healy, and Melville 2020) (**Fig. 4D**) shows clearly delineated clusters of fluorosequencing reads dominated by concordantly assigned reads, including separate clusters of the same peptide arising from distinct error modes, such as missing one Edman cycle (indicated by arrow (a)) or observation of a dud dye (indicated by arrow (b)), that can nonetheless still be correctly mapped to their corresponding peptide sequences.

### Targeted sequencing of an antigenic peptide

Finally, given these marked improvements in sequencing quality and the ability to match reads successfully to reference databases, we considered the goal of a clinical application and asked whether the enhanced workflow could detect HLA-I peptides from a small reference database of competing HLA-I peptides. The identification of tumor-associated peptide antigens in limited clinical biopsies poses a considerable obstacle for existing technologies, consequently hindering the advancement of novel therapies in immuno-oncology (Schumacher and Schreiber 2015; Vizcaíno et al. 2020).

We thus set up a pilot-study where we cultured ∼300 million mono-allelic B-cells (HLA A2603) and characterized the observed and potential HLA-I peptide repertoire through three orthogonal techniques (**Fig. 5A**). Despite the observed genomic variants and 3,237 predicted strong binding neoantigenic HLA peptides found through RNA sequencing, the vast majority of HLA-I peptides (1,189) identified by MS were distinct HLA-I peptides. Overall, fewer than 50% of the MS-observed peptides were predicted computationally to be strong HLA binders, possibly due to bias in current computational prediction algorithms (“The Problem with Neoantigen Prediction’’ 2017). We observed 4 putative peptides, identified through mass spectrometry and containing an alternative residue relative to the Hg38 reference human genome, serving as model neoantigens. We synthesized, fluorescently labeled, and sequenced one such peptide (*JSP308*, **Supplementary Table ST2**). Fluorescent radiometries indicated the correct step drop position for the Atto643 and Texas-red labeled residues (**Fig. 5B**). As above, we simulated fluorosequencing of a reference set of 37 HLA-I peptides (the set of MS-identified HLA-I peptides with more than two peptide spectral counts) including the input peptide, and trained a machine learning classifier to assign reads to the reference database peptides. Each experimental read was then assigned to a reference database peptide using this classifier, measuring classification accuracy and visualizing the reads by UMAP (**Fig. 5C**). As with the four peptide mixture, the target HLA-I peptide could in be clearly distinguished from the background set. We expect this trend to continue to improve with additional dye colors (allowing more residue types to be labeled) and other improvements, suggesting fluorosequencing may be applicable to the challenge of targeted identification of low abundance HLA-I peptides.

**Figure 5:**
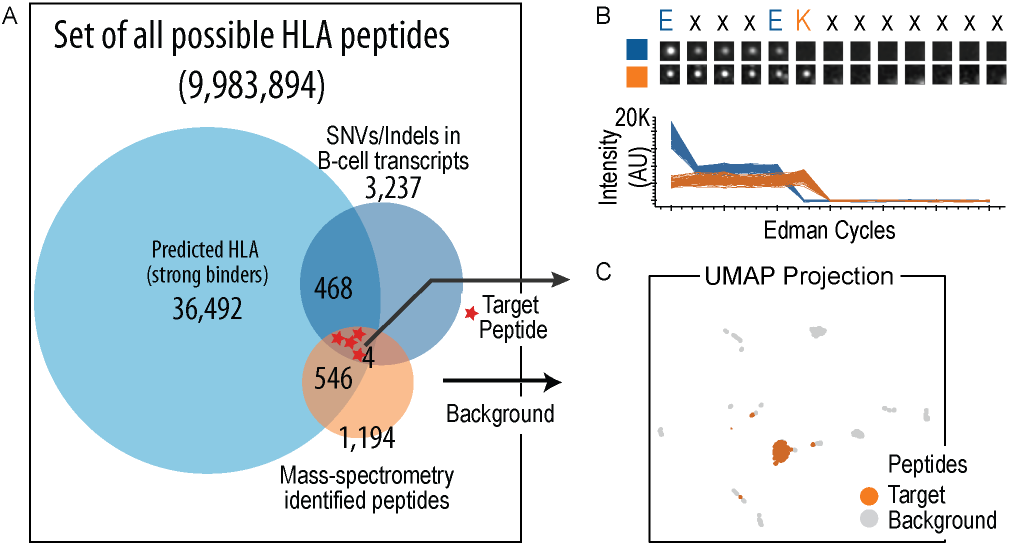
Sequencing of a target HLA-1 peptide demonstrates a potential application of fluorosequencing to address clinical need. (A) A pilot study was conducted on a mono-allelic B-cell line (HLA A2603) to compare potential neoantigenic HLA-1 peptides inferred from genomic and transcriptomic information with direct identification through mass spectrometry. The study revealed significant disparity of HLA-1 peptides, predicted through prediction algorithms and direct measurements. Out of 1194 peptides identified from mass spectrometry, only 546 were predicted to have strong affinity, and potentially four peptides with mutations were noted. Since the sensitivity requirement of detection of these peptides are typically high, we chose one of these peptides (JSP308) and labeled and fluorosequenced it. The experimental reads were classified against a reference database of the 37 well-observed HLA-1 peptides (observed with more than 2 peptide-spectral matches) from the larger set of 1,194 mass spectrometry-identified peptides. (B) Accurate sequencing of the peptide is illustrated by a representative image of one individual peptide molecule across Edman cycles (image series) and measured fluorescence intensities of 111 replicate peptide molecules (plotted values). (C) Despite the target peptide sequence having only three fluorophores and several similar peptides in the database (because HLA-I peptides bound by the same HLA allele share partial sequence identity, in this case at the peptide E1 position), 30% of experimental reads could still be correctly identified among those 676 reads classified with scores ≥ 0.7. This is illustrated visually in a UMAP projection, plotted as in Fig. 4D.

## DISCUSSION

After the initial proof-of-concept demonstration of the fluorosequencing technology, we made numerous improvements to the workflow to facilitate scaling the technology to identify multiple peptides in mixtures. Some of these improvements included selecting fluorophores that were stable across the Edman solvents and exhibited high brightness for single molecule TIRF experiments, optimizing Edman chemistry for greater reproducibility and efficiency, and modifying peptide-slide attachment through the azide-alkyne click reaction. During our experiments, we noticed that dye-dye interactions occurred between closely spaced fluorophores on the peptide. To overcome this issue, we installed the fluorophores on the peptide backbone using long, rigid linkers, called Promers. We designed the linkers to include a reactive chemical moiety, a flexible solubilizing PEG spacer, and a 30 unit proline polymer with a fluorophore on the other end. By spacing the dyes through these linkers, we minimized the quenching of similar fluorophores and FRET phenomena between different fluorophores. In addition, we created a new computational pipeline that increases the speed of image processing and scalability by inferring peptide identity in complex mixtures.

We also demonstrated our improvements to the fluorosequencing workflow by sequencing a mixture containing four synthetic peptides. After training our machine learning classifier on these peptides and 50 randomly generated decoy peptides, we were able to correctly classify 6% of the fluorosequencing reads with 99% precision, providing general support that the improvements to peptide labeling and workflow enables accurate peptide sequence determination. In a pilot experiment, we evaluated the ability to accurately identify a target antigenic peptide through a multi-omic study on an HLA A2603-expressing monoallelic B cell line. Using HLA prediction software, we found the set of expressed HLA peptides using genomic sequence and SNV comprising transcripts and compared it with those HLA-I peptides identified experimentally from the cell-line using tandem mass-spectrometry. We identified four peptides as a set of putative neoantigens. Fluorosequencing of one of HLA-I peptides against a reference background illustrates the potential and sensitivity for targeted clinical assays.

Computationally inferring peptide identity from fluorosequencing data has also suggested a workflow for *virtual fluorosequencing, i*.*e. in silico* modeling the entire sequencing process from proteins to expected fluorescence patterns for each possible generated peptide. The model is based on a rigorous characterization of the physicochemical processes and estimation of parameters, such as the probability of Edman failure, peptide detachment rate, chemical destruction rate etc (M. B. Smith et al. 2023). Virtual fluorosequencing can be used to guide experimental design by suggesting the protease and amino acids to label to yield the fluorescent pattern that best identifies the peptide or protein of interest, as well as to identify the most sensitive parameters that, when improved, would lead to better peptide identifications. By guiding experimental design, virtual fluorosequencing provides a valuable tool for improving the reliability of fluorosequencing.

The successful completion of the end-to-end process for fluorosequencing of peptide mixtures is a major milestone for the technology. In this study, we demonstrated several of its key features, including high sensitivity and the generation of rich quantitative data, that make fluorosequencing a promising tool for clinical applications requiring accurate peptide and protein measurements from low sample amounts.

## Supporting information

All Supplemental Tables

## ACKNOWLEDGEMENTS

We gratefully acknowledge the UT Chemistry Mass Spectrometry Facility for their assistance and service in analyzing samples. MALDI and MS/MS data collection was supported by CPRIT grant RP110782 to Dr. Maria Person. We thank Dr. Mariah Austin for designing the figures. We would like to thank Drs. Luke Lavis (Janelia Research), Rajendra Singh (Goryo Chemicals), Martin Schnermann (NIH), Taki Masayasu (Nagoya University), Yamaguchi Shigehiro (Nagoya University), and Ramesh Jasti (University of Oregon) for generously providing us with either fluorophores or relevant information about them for our dye screen. Research at the University of Texas was supported by a Sponsored Research Agreement from Erisyon to E.M.M. and E.V.A.; E.M.M. acknowledges additional support from the Welch Foundation (F-1515) and NIH (R35 GM122480). E.V.A. acknowledges support from the Welch Regents Chair (F-0046).

## AUTHOR CONTRIBUTIONS

All authors assisted in design, execution and interpretation of the data. JS wrote the manuscript with the assistance and inputs from other co-authors. All authors edited the manuscript.

## DATA AVAILABILITY

Raw fluorescence images, processed fluorosequencing data, results of HLA peptide prediction, and characterization data for all the peptides presented in the manuscript have been made available on Zenodo at doi:10.5281/zenodo.7958185. Mass spectrometry proteomics data have been deposited in the proteomeXchange database under accession number PXD042439.

## CODE AVAILABILITY

All code used in the paper has been made freely available under a permissive MIT license at https://github.com/marcottelab/robust-fluorosequencing-plaster.git. Note that operating the code requires implementing back-end Amazon Web Services.

## MATERIALS AND METHODS

Our materials and methods are summarized concisely in this section. Additional details about the microscope and fluidics, fluorescent dye screen, and computational workflow are provided in the **Supplementary Notes**.

### Purification methods

#### Using analytical column chromatography

We used an HPLC system (1100C, Agilent) with an analytical column (Kinetex 5 μm XB-C18, 250 × 4.6 mm, 100 Å) operating at a flow rate of 1 mL/min and a 30 min gradient (5-95% Acetonitrile/water with 0.1% Formic acid) to purify samples with <1mg peptide or fluorescently labeled peptide. The different fractions were collected using the attached fraction collector and the species were analyzed using LCMS. The purified peptide fraction was then dried down using speed-vac (Eppendorf) and solubilized in 50% Acetonitrile/water for further characterization.

#### Using preparative or semi-preparative scale HPLC

We used a semi-preparative or preparative HPLC system for samples containing larger amounts of peptides (>1mg). The preparative scale purification involved using an Agilent Zorbax column (4.6 × 250 mm) on an HPLC system (Shimadzu model) operating at a flow rate of 10 mL/min, and eluting with a 90-minute gradient of 5–95% acetonitrile (0.1% Formic acid). For the semi-preparative scale purification, we purified the product using HPLC (Shimadzu) with a semi-prep column (Hichrom C8, 5 micron, 10cm x 10 mm, 150 Å) operating at a 5 mL/min flow rate and an elution gradient of 5-95% Acetonitrile (0.1% Formic acid) over 60 minutes. We analyzed the fractions using mass spectrometry and pooled the fractions containing the product. We then reduced their volume using a roto-vap (IKA, RV10) and lyophilized the samples (VirTis SP Scientific, BTP-8ZLOOW) prior to characterization.

#### SDS PAGE gel purification

For peptides labeled with fluorescent promers, we performed standard SDS PAGE electrophoresis on a 16.5% Tris/Tricine SDS PAGE gel, using vendor detailed protocol (Biorad, Cat# 1610739, #4563065, #1610744). After washing the gel and confirming the fluorescent bands using a gel imaging station (Amersham Imager 600 gel dock), we cut the bands of interest using a razor blade. The excised pieces were crushed and submerged in 50% vv of Acetonitrile/water in a microcentrifuge tube. We sonicated them for 5 minutes and heated them at 60C for 30 minutes to extract the peptides from the gel.

### Synthesis of peptides

All peptides were either custom synthesized from Genscript (NJ, USA) or synthesized in-house using a standard automated solid-phase peptide synthesizer (Liberty Blue microwave peptide synthesizer; CEM Corporation). If synthesized in-house, we cleaved the peptides from beads with standard TFA cleavage cocktail (comprising 95% TFA, 2.5% water and 2.5% Triisopropylsilane (Sigma, Cat #233781) for 2-4 hours at room temperature, followed by ether precipitation. After that, we purified the crude precipitate by preparative scale HPLC. We characterized the peptides synthesized by LCMS.

### Synthesis of Promers

The amino acid sequence of the Promers, as read from N-terminus to C-terminus, is Fmoc-G-PEG2-G-P_30_-K(boc)-CONH2. We used standard solid-phase Fmoc synthesis, and performed double coupling of proline residues after the first 20 amino acid synthesis to build the rest of the chain. We removed the terminal Fmoc group and reacted the resin with 5 equivalents (eq) of DBCO-NHS (dibenzocyclooctyne-N-hydroxysuccinimidyl ester) in dry DMF, containing 2.0 eq of triethylamine for 2 hours at 37°C to functionalize the polypeptide with DBCO. After washing and cleavage and HPLC purification of the DBCO-functionalized polypeptide, we labeled the ε-lysine on the polypeptide with 1.2 eq of succinimidyl ester derivatized fluorophores (Atto643, JF549, or Alexa555) by incubating it in 1:1 HEPES (0.1M, pH 8.5), acetonitrile buffer for 2h at 37°C. We HPLC purified the fluorescent Promer-DBCO product using a semi-preparative column.

### Fluorescent labeling of peptides

#### Direct fluorescent labeling

In general, we coupled 0.9 eq of NHS functionalized dye with ∼100 nmoles of peptides, per lysine, by incubating the mixture in 100 μL of 1:1 vv Acetonitrile/HEPES buffer (pH 8.5, 0.1M) at room temperature for 2 hours. After the reaction, we purified the peptide using an analytical column. For peptides containing both lysine and cysteine residues (Peptides: *JSP129, JSP150* and *JSP157*) and requiring two different fluorophores, we first labeled the cysteine residue by incubating the peptide with 1.2 eq of Atto647N-Iodoacetamide (Atto-tec, Cat# AD 647N-111) in 100 μL of sodium phosphate buffer (pH 7.5; 0.1M) and incubating for 2 hours at room temperature. We then adjusted the pH to 8.5 by adding 50 μL of 1M HEPES (pH 8.5) and added 1.2 eq of Fluorophore-NHS. This was incubated overnight at room temperature, followed by HPLC purification using the analytical column.

#### Labeling with fluorescent Promers

Across different peptides, we incubated 100 nmoles of peptides containing azidolysine directly with 2 eq of DBCO functionalized dye Promers in 100 μL of 1:1 vv Acetonitrile/HEPES buffer (pH 8.5, 0.1M) at room temperature for 16h. After the reaction, we purified the peptide using the analytical HPLC. If peptides required two different fluorophores, we used a two-step method by first labeling the azidolysine residue with the Promer, followed by functionalizing the lysine residue with NHS-PEG4-Azide (Broadpharm, Cat #BP-20518). We then purified the resulting peptide using the analytical column (described earlier) and labeled the second fluorophore using the same DBCO azide chemistry. Finally, we purified the two-color peptides using either the analytical HPLC column or SDS PAGE gel.

### Isolating and identifying HLA peptides using tandem mass-spectrometry

We purchased the monoallelic B-cell line (B721.221), with HLA allele A*2603, directly from the International Histocompatibility Working group (Seattle, Washington). We cultured the cells and isolated HLA peptides from approximately 300 million cells as previously described (Abelin et al. 2017). We identified the isolated HLA peptides using an LC-coupled tandem mass-spectrometer (ThermoFisher, Orbitrap Fusion Lumos) and a reference dataset of a human proteome (Swissprot), applying settings from the literature for analyzing HLA peptides (Bassani-Sternberg et al. 2015) and ProteomeDiscoverer 2.3 software (Thermofisher). Raw data has been deposited to ProteomeXchange Consortium via the MassIVE partner repository with data set identifier PXD042439.

### Predicting HLA antigenic peptide from genomic and transcriptomic data

We obtained the RNA sequencing data for the B cell-line (expressing HLA-A2603 allele) from a publicly available dataset (Abelin et al. 2017). We performed RNAseq alignment using STAR tool (Dobin et al. 2013) and single nucleotide polymorphism analysis by comparing the aligned transcriptome to the standard human genome (Genome Reference Consortium Human Build 38) using GATK best practice pipeline (https://gatk.broadinstitute.org/hc/en-us). We manually translated 50 base pairs of 5’ and 3’ from the identified functional single nucleotide variant (Wang, Li, and Hakonarson 2010) to peptide sequences. We then predicted the presented HLA peptides using netMHCPan 3.0 software (Lundegaard et al. 2008). We subsequently filtered the peptides for strong binding to the HLA allele A*2603. All associated data for antigen peptide prediction is uploaded to Zenodo (doi:10.5281/zenodo.7958185)

### Characterization of peptides

#### LC/MS

We characterized the peptide samples using Liquid chromatography mass spectrometry (LCMS) with an Agilent 1260 system equipped with Single Quadrupole LC/MS (6120B) and a 5 μm C18 column (Agilent Zorbax Eclipse Plus, PN 959746-902). We injected the samples and subjected them to an elution gradient of 5-95% aqueous (0.1% FA) over 12 minutes with a co-eluent of Acetonitrile. We used Agilent Chemstation (ver # Rev C.01.10 [287]) to analyze the data.

#### MALDI

We characterized peptides containing Promers with a mass range of 3K-20K using MALDI TOF (Autoflex max, Bruker). We spotted 1 μL of solubilized sample onto a clean MALDI Target plate (MTP 384 target plate polished steel, Bruker) and mixed it with 1-2 μL of 40 mg/mL DHB (Thermofisher, Cat #90033) in 70% Acetonitrile and 0.1% Trifluoroacetic acid. After drying, we analyzed the sample in reflective mode at a laser power of 60-90%. We used Autoflex analysis software (ver #3.4) to analyze the data.

### Fluorescence reading using plate reader

Fluorescent peptides were diluted into methanol to an ∼10 μM concentration and fluorescence measured using the fluorescence plate reader (Synergy H1 microplate reader, Biotek-Agilent). The samples are excited at 500 nm and emission measured from 520-700 nm in increments of 10 nm. No gain setting was used.

### Silane functionalization of glass slides

We cleaned 40 mm glass cover slides (Bioptechs, Cat 40-1313-03192) for 10 minutes on each side using an UVO cleaner (Jelight, Model 18). After cleaning, we placed the slides vertically in a Teflon slide rack (custom-made). We then pipetted 100 μL of 3-azidopropyltriethoxysilane (Gelest, SIA0777, CAS# 83315-69-9) into the lid of a Teflon Reaction Vessel (Alpha Nanotech Inc) and placed both the slide rack and the cap in a Pyrex desiccator chamber, which we had preheated to 80C. We attached the valve of the desiccator to a vacuum pump and drew a vacuum until the pump stabilized at approximately 0.08 MPa. We then placed the desiccator in an 80C oven and allowed it to sit for 16 hours. We stored the silane-functionalized slides in vacuum-sealed bags at 4C until use.

### Peptide immobilization

Peptides (containing alkyne) were covalently coupled to the coverslip surface via copper-catalyzed click chemistry between the alkyne-modified C-terminal AA residue and the azido silane. A fresh solution of 2 mM copper sulfate, 1 mM tris(3-hydroxypropyltriazolylmethyl)amine (Sigma, Cat # 762342), 20 mM HEPES (pH 8.0), and 5 mM sodium ascorbate with fluorescently labeled angiotensin was incubated for 30 min at room temperature on the coverslip, washed with water to remove unbound peptides, and dried under a nitrogen gas stream.

### Total Internal Reflection Fluorescence (TIRF) Microscopy

Two similar Nikon Ti microscopes, equipped with a CFI Apo 60X/1.49NA oil-immersion objective lens and a 1.5× tube lens, a motorized stage, a sCMOS camera, and a laser excitation were used for all the experiments. Details of these parts and the fluorescent channel configurations are provided in the **Supplementary Notes**.

### Automated fluidics for performing Edman sequencing chemistry

#### Fluidic setup

We automate the pumping of different solvents using a syringe pump (Tecan Cavro, Model# 20738291) (3 way valve configuration) and a 10-port multi-position valve system (Valco Instruments, Model# EUHB), as described in the earlier publication (Swaminathan et al. 2019). We maintain the sample temperature at 40C (for System A) and 50C (for System B) by heating both the perfusion chamber and microscope objective for Edman sequencing experiments. We control solvent exchanges in the fluidic device using in-house Python scripts and coordinate them with image acquisition via custom macros in the Nikon Elements software package. We connect the reagents/solvents to the different valves and detail the steps for performing Edman chemistry in **Supplementary Table ST7**.

### Signal processing

We extracted raw sequencing reads from the time series micrographs as detailed in the **Supplemental Notes** and implemented in the Python package *sigproc_v2*, available from https://github.com/marcottelab/robust-fluorosequencing-plaster. Briefly, images capturing the same field of view were corrected for variation in signal intensity by regional illumination balancing and bandpass filtering of both background and over-saturated pixels, aligned across cycles to account for stage movement, and peaks identified by convolving with the point spread function (PSF). From radiometry on each peak, we calculated intensity parameters for each peak across Edman cycles to create a raw fluorosequencing read for each candidate peptide. Fields were filtered on the basis of image anomalies or poor alignments, and individual reads were rejected based on their fit to the PSF, with poor fits suggesting the presence of more than one molecule. Filters are detailed in **Supplementary Table ST8** and additional filters are described in the **Supplemental Notes**.

### Scoring co-localization

For each raw fluorosequencing read (acquired, extracted, and filtered as described above) the intensities were normalized to the mean one-count intensity (μ) and plotted by channel pairs as scatter plots; **Fig. 3D** shows an example. Signal above the dark thresholds (determined at 3 σ above the background mean) in both channels were considered colocalized and were calculated as the percentage of the total reads plotted.

### Measuring FRET efficiency

For FRET analysis, the donor (lower wavelength) and acceptor (higher wavelength) dyes were imaged as separate channels using their respective laser and filters. We additionally defined a “FRET channel” using the donor’s excitation laser and filter and the acceptor’s emission filter. Details of the System A setup can be found in the **Supplementary Notes**. Data was acquired in all channels and the reads were calculated and filtered as described above. These reads were then used to calculate the FRET efficiency as in (Hellenkamp et al. 2018; Zal and Gascoigne 2004). To simplify the analysis, we also filtered for peptides with signals in all channels above the contamination background.

We calculated FRET efficiency (*E*) for each read using **Eq. 1**, where *I*_*F*_, *I*_*D*_, and *I*_*A*_ represent the fluorescence intensities of the FRET, donor, and acceptor channel reads, respectively.

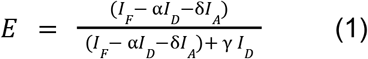

This formula requires the determination of experimental coefficients to account for the modifications of the intensities inherent in this type of data acquisition. The first of these coefficients (*α* and ***δ***) account for signal leakage into the channels that occurs from overlapping excitation or emission (crosstalk). To calculate these, reads from a control experiment with single labeled peptides were plotted in the same manner as the colocalization analysis. The slope of the correlation between the signals in each channel pair was used as the crosstalk coefficient giving a value of 0.20 for the donor to FRET channel (*α*) and 0.03 for the acceptor to FRET channel (***δ***) for the peptides in **Supplementary Fig. SF8B**. The last coefficients (*γ* and *β*) is to normalize effective fluorescence quantum yields, as these experiments are on purified peptides, we used the ‘single-species’ method described in Hellenkamp et al. giving value of for *γ* and *β* asn 1.2 and 2.9 for the peptides shown. To confirm the these coefficients we calculated the stoichiometry (*S*) for each peak using **Eq. 2**, and confirmed that the mean stoichiometry was the expected 0.5, for these peptides having one donor and one acceptor fluorophore.

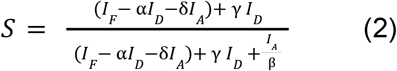

### Machine learning classifier

To generate synthetic training and testing data typical of fluorosequencing experiments, we performed Monte Carlo simulations as in (M. B. Smith, Simpson, and Marcotte 2022), explicitly modeling the Edman failure rate, peptide detachment rate, and N-terminal blocking rate, as well as a number of fluorophore-specific parameters, including each dye’s average fluorescence intensity, standard deviation of intensity, standard deviation of background intensity, missing (“dud”) dye rate, and dye destruction rate (a combination of chemical destruction and photobleaching rates, the latter of which is small in the conditions used for imaging). Methods for estimating these parameter values are detailed in the **Supplementary Notes** (with additional methods available in (M. B. Smith et al. 2023) and all parameters are provided in **Supplementary Table ST4**. These synthetic reads were expressed as double-precision floating point vectors comprising (simulated) fluorescence intensities for each Edman cycle and fluorescent channel. By considering these reads as feature vectors, we trained a random forest classifier to assign reads to source peptides, as implemented with scikit-learn and default settings.

