## Extended Data Figures and Supplemental Notes for "Robust and scalable single-molecule protein sequencing with fluorosequencing"

**SFIG 1A: Atto643 and TexasRed dyes were selected as current optimal dyes for fluorosequencing through controlled sets of single molecule experiments.** We downed selected dyes - Atto643 and TexasRed through estimating and comparing parameters from a controlled set of experiments. (i) Comparing dye-destruction rate through cycles of Edman chemistry on acetylated peptides (JSP260 and JSP288) carrying their respective fluorophores, show the rates are 5.6% and 2% per cycle (ii) Comparison of photobleaching rates between these peptides shows that less than 1.1% and 19% of Atto643 and Texas Red dyes photobleach in 15 imaging cycles. (iii) mean intensity (mu) and the spread (sigma) of Atto643 dye is 11729.35 and 0.22 and TexasRed is 4970.85 and 0.19 AU respectively.

**SFIG 1B:: Solvent stable fluorophores were selected to span the visible spectra.**

To enable fluorosequencing, we need multiple fluorophores that can be distinguishable across the visible spectra. Through screening of 70 dyes for solvent stability, we have identified four different fluorophores, Atto425, Atto495, TexasRed, and Atto643, for our microscope imaging setup (see methods). The excitation and emission spectra for each of these fluorophores are shown.

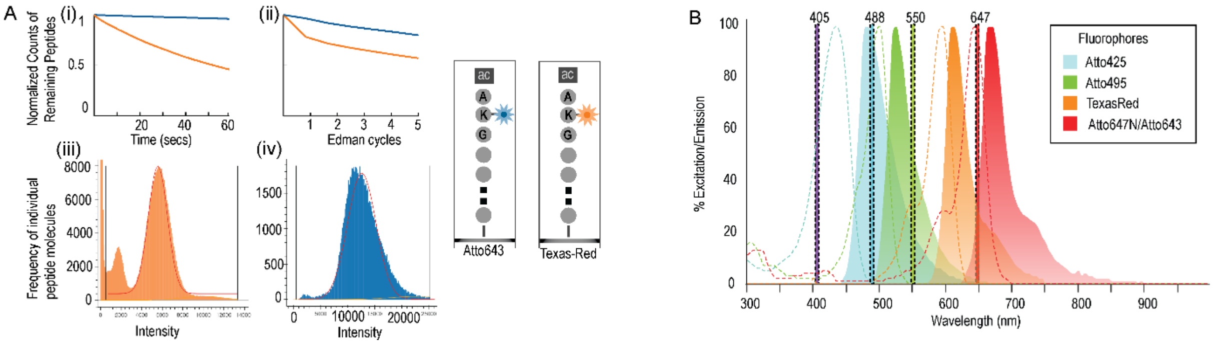

**SFIG 2: Optimization of coupling solvent and time for cleavage chemistry increased Edman efficiency to > 95% across a range of different peptides. (Panel A)** The normalized counts of fluorescently labeled amino acid cleaved at the correct cycle, increased with increasing time of TFA incubation with maximum cleavage rate observed with 8 min trifluoroacetic acid incubation time. **(Panel B)** The addition of N-methylmorpholine (x mM) into the PITC coupling solution, increased the Edman efficiency by 12% (depicted as drop percentage of fluorescent tracks at 2nd position) for peptide JSP127.

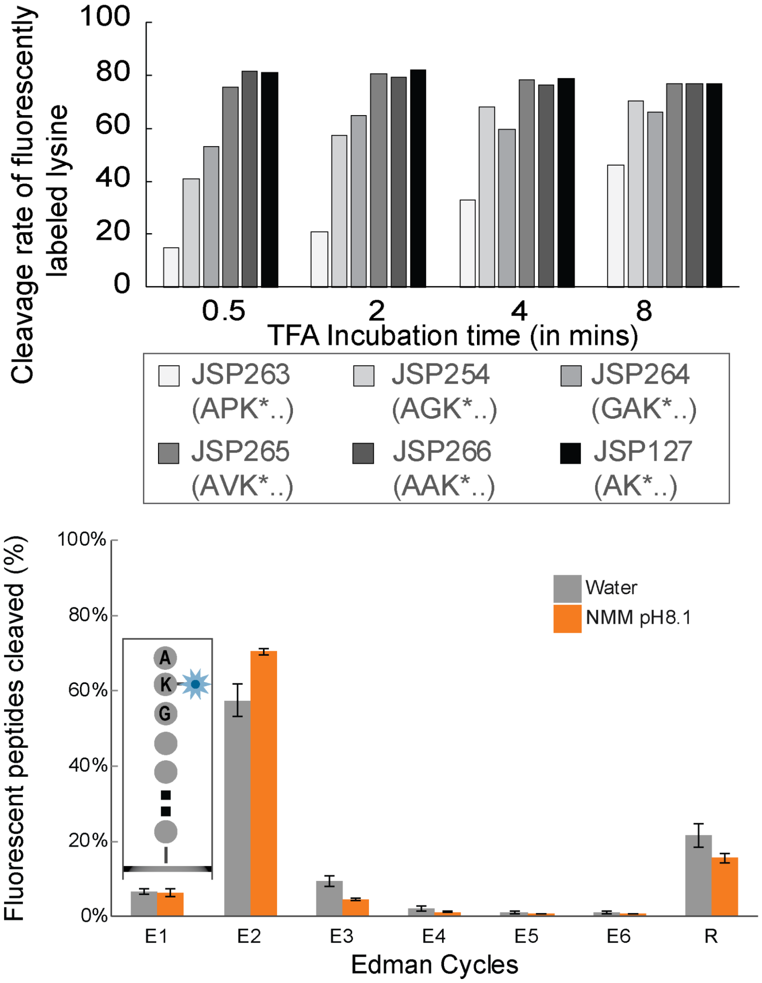

**SFIG 3: The position of prolines with respect to the labeled amino acid effects the efficiency of Edman degradation.** We observed that the efficiency at which the Atto643 labeled lysine residue is cleaved is affected by the presence of proline residues by an average of 9.5%, when proline residues were located N-terminal to the fluorescently labeled lysine residue. Fluorosequencing for two sets of similar peptides were compared, differing in the position of proline residue at the 2nd (panel A; peptides - *JSP286, JSP263*) or 3rd position (panel B; Peptides - *JSP285, JSP287*), and the decrease in efficiency was found to be similar.

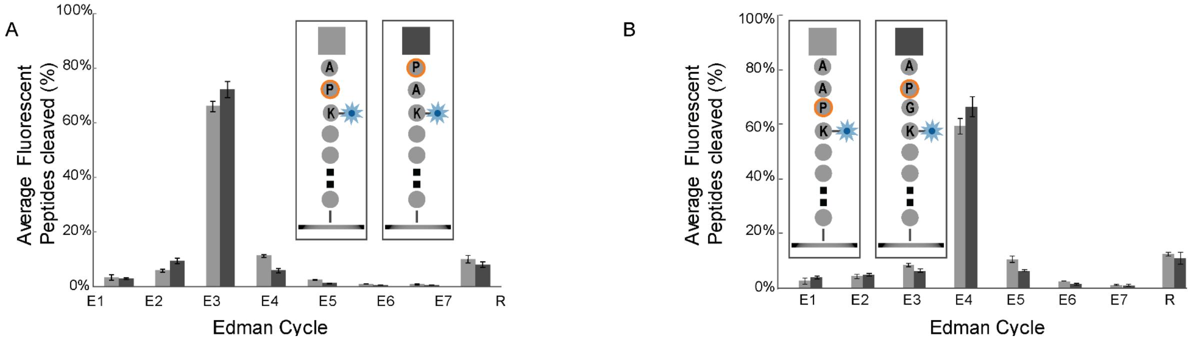

**SFIG 4: Vapor deposition method for silanization of glass slides reduces fluorescent contamination across the different imaging channels.** (A) Image setup for vapor deposition of 3-azidopropylsilane (see methods) is shown. (B) The fluorescent images of the slide post functionalization shows extremely low counts of fluorescent contaminants across the four imaging channels (445, 480, 532 and 561 channel; see methods for optical setup). The values represent the number of peaks/field for each channel. The peptides in the 640 channel (not shown) contains a dye-labeled peptide and used to focus the slide. Scalebar represents 10μm

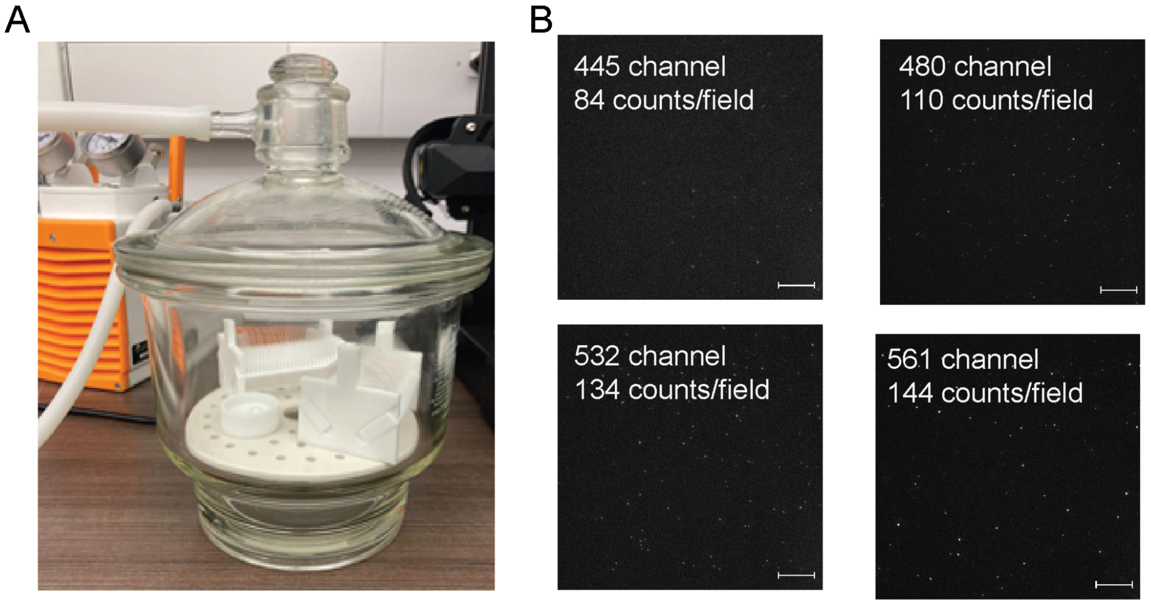

**SFIG 5: Illustration of the computational workflow for inferring peptide sequences from raw fluorosequencing data**. Briefly, the workflow comprises of two major parts - (a) building of a machine learning classifier: Using the input peptide set and experimentally determined fluorosequencing parameters, namely, Edman efficiency, photobleaching rates, dye destruction rates, dud dye rates and dye intensity distributions, we simulate possible fluorosequencing reads. With the knowledge of the source peptide, we train and test using random forest to build a classifier. (b) Image processing: the raw images obtained for 1000s of images across different fluorescent channels and Edman cycles are collected, aligned, filtered and fluorescence intensity reads obtained. Each fluorescent track is then classified to an input peptide with a score. Applying a score threshold, we collate the counts of individual peptides present in the input sample. Details of the protocol is given in methods section and pseudocode.

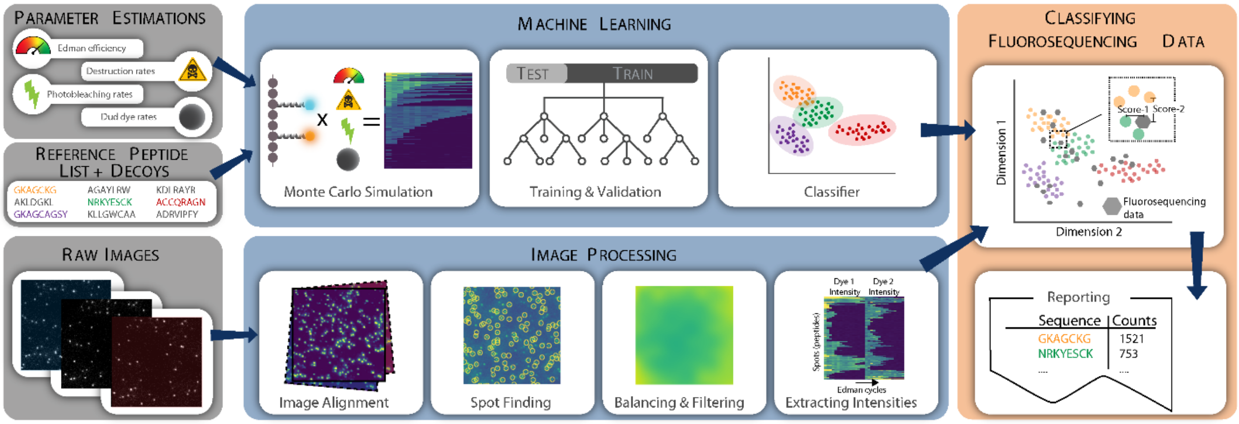

**SFIG 6**: **Quenching of fluorophores labeled on the same peptide was observed**. By measuring the intensity distributions for single, two, three and four fluorophores and attempting to fit them through an additive gaussian distribution, we observe that there is significant dye-dye interactions or quenching of fluorescent signal between the fluorophores. The raw intensities for the four different peptides are shown in panel A-D, with an attempted overlay of the predicted intensity distribution based on the gaussian fit parameters for the single dye.

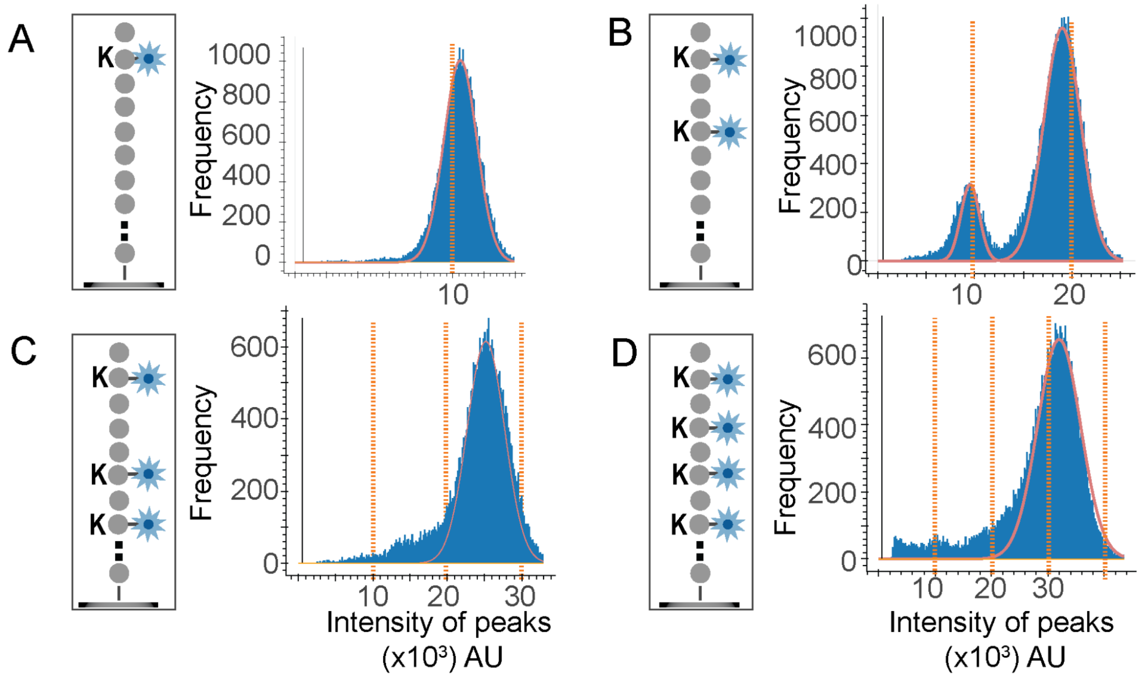

**SFIG 7: FRET is observed across wide range of donor fluorophores**

**SFIG 7A-B: FRET phenomena is observed in peptides containing JF549 and Atto647N.** (A) Overlay and offset images of the peptides across three channels - (1) 561, (2) "FRET" and (3) 561 channel indicates the missing signal in the 561 or donor channel. (b) Recovery of the counts of the 561 channel after photobleaching of the dyes in the 647 channel can be seen through the raw images of the donor and the acceptor channels before and after photobleaching

**SFIG 7C-D: FRET phenomena is also observed across multiple combinations of dye-pairs - Alexa488/Atto647N and JF525/Atto647N.** Despite the minimal overlap between the emission spectra of the Alexa488 and JF525 dyes with the Atto647N dyes with estimated FRET efficiency of 32.2% and 37% (as calculated using the online FRET calculator - https://www.fpbase.org/fret/), we observe significantly high FRET signal.

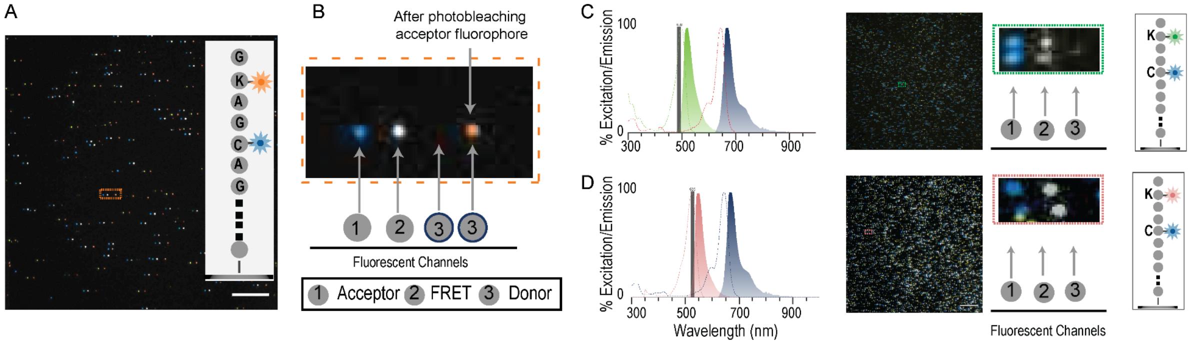

**SFIG 8: FRET is mitigated between fluorophores on peptides, when attached through polyproline linkers**

**(Panel A SFIG 8A):** Donor signal of tetramethylrhodamine, recovers when spaced away from Atto647N dye using rigid polyproline linker. We excited the donor fluorophore (Tetramethylrhodamine) on two peptides (depicted in the legend in the left panel) using a 500 nm monochromator. We recorded the emission signal from 525-700 nm and found that the tetramethylrhodamine dye spectrum was absent for the shorter Pro(3) peptide while present in the Pro(14) (Peptide-JSP168).

**SFIG 8B: Increased polyproline linker length to 30 units decreased FRET efficiency to <10% on the single molecule imaging system.** We performed single molecule imaging on three peptides, JSP212, JSP213, and JSP214, with different constructions of donor fluorophore (Janelia fluor 549) and acceptor fluorophore (Atto643). The left panel illustrates the fluorophore constructions, indicating the presence or absence of a polyproline linker (shown as a helix) on the three peptides. (A)The scatter plot of the intensity of peptides across the 560 and 640 channel is shown for each of the three peptides. The peptide with no Pro(30) linker (top figure) had only <5% of colocalized spots, while high colocalization (67%) was observed when fluorophores were constructed with a Pro(30) linker (bottom). (B) The FRET efficiency across the three peptides for each of the individual peptide measurements is shown. The stoichiometry value for every individual peptide measurement is the ratio of donor and acceptor fluorophore after normalization of intensity and cross-talk across the channels. The spacing of fluorophores through the construction of a Pro(30) linker reduced FRET efficiency to less than 10% (shown in the bottom figure).

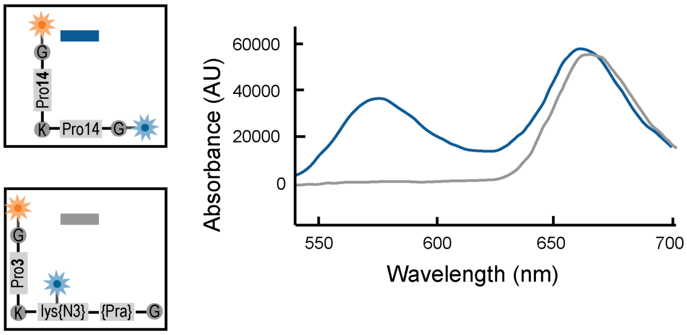


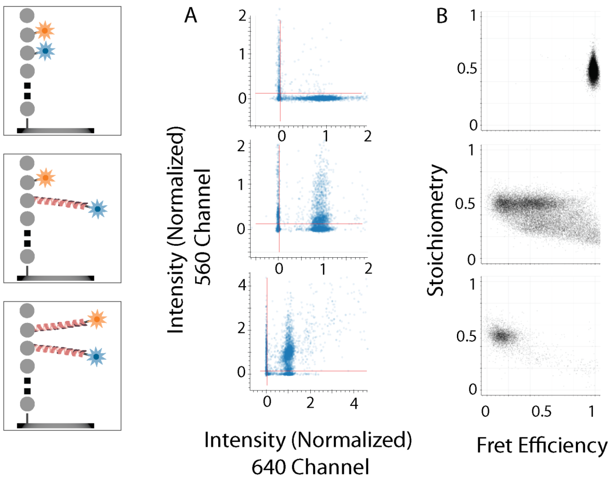

**SFIG 9: Edman degradation occurs at similar rates for peptides containing fluorescent Promer.** There was no significant difference in Edman degradation efficiency observed between peptides with fluorophores constructed with and without promers (JSP263, JSP274), as indicated by the average loss of 62% of fluorescent peptides at the 3rd position.

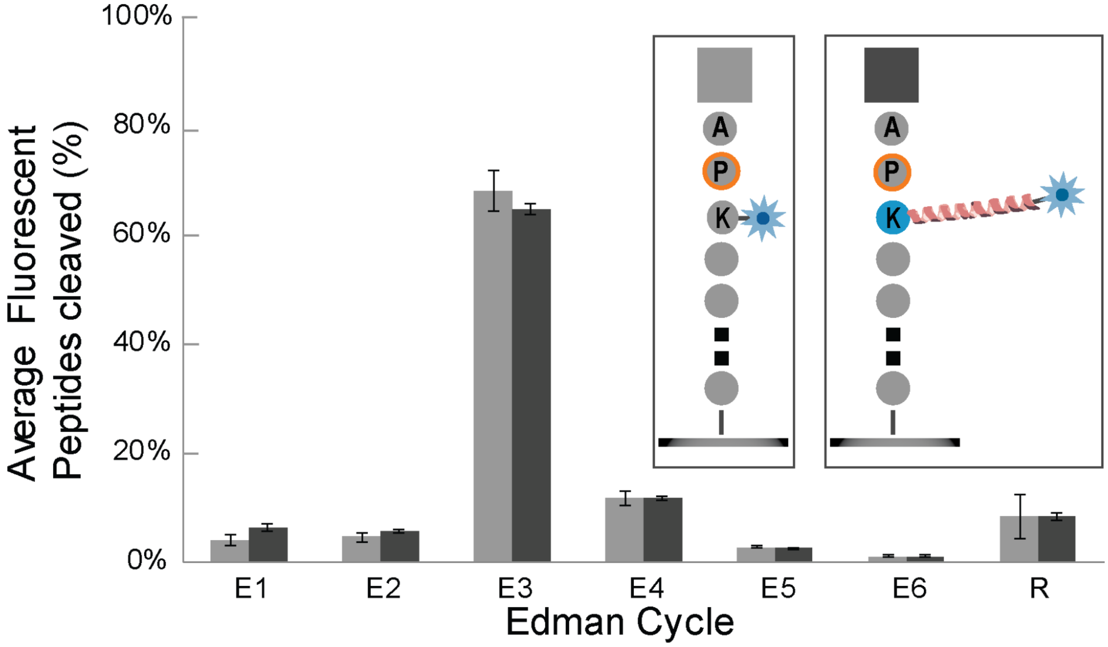

**SFIG 10: Score distribution between decoy and input peptide set.** The results of the experiment scored 49,480 fluorescent tracks to the most likely peptide, either to the 4 peptides present in the input set or the decoy peptides (see methods). Panel A presents a histogram showing the count of peptides for various classification scores. It is clear from this data that higher scores correspond to peptides from the input set, which are represented in blue. On the other hand, the decoy peptides are not highlighted. In Panel B, the peptides that have been classified are evaluated on a precision/recall curve. The data suggests that 8% of the peptides (in terms of recall) had a very high precision of 99%.

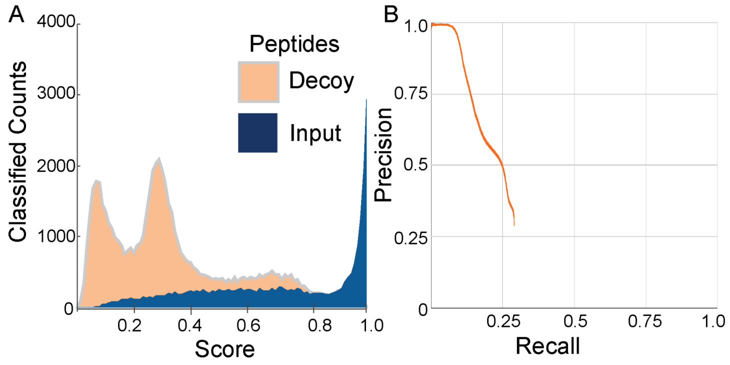

### ADDITIONAL DESCRIPTIONS OF METHODS AND MATERIALS

#### SDS Page gel purification

For peptides labeled with fluorescent Promers, we mixed them with a Tricine Sample Buffer (BioRad, Cat#1610739) and loaded them onto a 16.5% Tris/Tricine SDS PAGE gels (BioRad, Cat#4563065) with Tris/Tricine running buffer (BioRad, Cat#1610744) while excluding traditional reduction and heating during sample preparation. We ran gel electrophoresis (Biorad, Cat#4006213) on the loaded sample, until loading dye ran off the gel. After washing the gel, we imaged the gels using a gel imaging station (Amersham Imager 600 gel dock) in the 530 and 630 nm fluorescent channels. We cut out bands of interest using a razor blade, washed the excised pieces with water, and then crushed and submerged them in a 50% vv of Acetonitrile/water in a microcentrifuge tube. We sonicated them for 5 minutes and heated them at 60C for 30 minutes to extract the peptides from the gel. Then, we removed the supernatant and used a C18 ziptip (Thermofisher) to desalt and purify the peptides. We characterized the excised peptides using LC-MS or MALDI. We found that Promer labeled peptides had a different migration speed than protein standards; a 10kDa peptide with Promers had a similar retention time as a 25kDa Protein standard (Precision Plus Protein Dual Xtra Standard, Cat#1610377).

#### Detailed synthesis of Promers

Using an automated peptide synthesizer (Liberty Blue microwave peptide synthesizer, CEM Corporation) to synthesize the polymer backbone of the Promers. We prepared fresh stock solutions of each amino acid building block (Fmoc-glycine, Fmoc-proline, Fmoc-lysine(boc), and Fmoc-PEG2) at a concentration of 0.2M, as well as coupling reagents (1M of Oxyma base and 1M of DIC). After coupling the first 20 monomers on the resin, we performed double coupling of Fmoc-glycine, Fmoc-proline, and Fmoc-Peg2. Then, we removed the terminal Fmoc group and reacted the resin with 5eq of DBCO-NHS in dry DMF, containing 2.0 eq of Triethylamine for 2 hours at 37C to functionalize the polypeptide with DBCO. Next, we washed and cleaved the DBCO-functionalized polypeptide from the resin with an acidic cocktail consisting of 50% TFA, 45%DCM, 2.5% Triisopropylsilane, and 2.5% water for 2h at room temperature. We dried the cleavage cocktail via $N_2$ gas until ≤5% of the initial volume remained, then added 10:1 vv of cold ether to precipitate the peptides. After decanting off the ether, we re-solubilized the resulting solid product in 50% Acetonitrile/Water for purification. We purified the DBCO-polypeptide using HPLC with a semi-prep column (Hichrom C8, 5 microns, 10cm x 10 mm, 150 Å) operating at a 5mL/min flow rate and an elution gradient of 5-95% Acetonitrile (0.1% Formic acid) over 60 minutes. Then, we labeled the Finally, we purified the labeled peptides using HPLC and the same semi-prep column as described earlier. We provide representative LCMS traces and MALDI characterization for the Promers used in this publication in the supplementary information.

#### Synthesis of peptides with N-terminal branched proline polymer

We synthesized a peptide with the sequence Boc-Lys[fmoc]-Gly-azLys-Gly-Pra-Gly-Resin on Tentagel Rink Amide Resin by using boc-lysine(fmoc) to enable the synthesis of variable length proline backbones from the lysine side chain. After synthesizing the proline polymer, we installed a terminal glycine residue. Then, we labeled the N-termini on the branched glycine with either TMR-NHS or JF549-NHS dye (1.2eq) on the resin by incubating it in DMF and 2eq of Triethylamine for 2h at room temperature. After cleaving the peptide and purifying it using preparative scale HPLC, we labeled the azidolysine on the peptide with 1.2 eq of DBCO-Peg4-Atto647N (custom synthesized by Atto-tec) by incubating the mixture overnight at room temperature.

**Fluorophore selection through solvent stability screen.**

We obtained the fluorophores commercially or obtained them through collaborators (see supplementary sheet 3 for source). We screened 70 fluorophores to identify those most resistant to the Edman solvents by covalently attaching the dyes to Tentagel beads (Chem-Impex International, 04773) and measuring their fluorescence after a 24-h incubation with TFA, pyridine/PITC (9:1 vv), Methanol and Piperidine at 40 °C. Nonspecifically bound fluorophores were removed by repeated washing with dimethylformamide (DMF), dichloromethane, and methanol. These beads labeled with fluorescent dyes were suspended in 100 µL of phosphate-buffered saline (PBS, pH 7.2) in a 96 well plate. We captured the fluorescent bead images across multiple channels, using an Epi-microscope and calculated the change in fluorescent intensity, compared to the methanol control. Custom script was used to measure the bead fluorescence from the images.

Epi-microscope (Nikon Eclipse TE2000-E inverted microscope) used was equipped with an Apo 60×/NA 0.95 objective, Cascade II 512 camera (Photometrics), a Lambda LS Xenon light source and a Lambda 10-3 filter-wheel control (Sutter Instrument), and a motorized stage (Prior Scientific), all operated via Nikon NIS Elements Imaging Software. Images were acquired at one frame per second through a 89000ET filter set (Chroma Technology) with channels 'DAPI' (excitation 350/50, emission 455/50), 'FITC' (excitation 490/20, emission 525/36) 'TRITC' (excitation 555/25, emission 605/52), and 'Cy5' (excitation 645/30 emission 705/72).

**Total Internal Reflection Fluorescence (TIRF) Microscopy**

Single-molecule TIRF microscopy experiments were performed on two different Nikon systems, detailed below:

**System A**

Nikon Ti-E inverted microscope equipped with a CFI Apo 60X/1.49NA oil-immersion objective lens and a 1.5X tube lens, a motorized stage (TI2-S-HW, Nikon Inc Scientific), an 1022x1022 pixel sCMOS detector (pco.edge, PCO),and a LUNF-XL (Nikon) laser including 561 and 647 nm lasers and filter cube containing 405/488/561/638 quad dichroic and barrier filters, an emission filter wheel with band pass filters detailed below (all filters, Chroma). Each image represents a 72 µm × 72 µm square region of the sample. The different channels can now be considered as a combination of incident laser wavelength and the corresponding bandpass filter. The “561 channel” consists of excitation with the 561 nm laser (9.5 mW, 50%) through quad dichroic and emitted signal collected through emission filter EM-603/30. The “640 channel” consists of excitation with the 640 nm laser (2.5 mW, 10%) and collected through quad dichroic and EM-705/72 emission filters. The “FRET channel” consists of excitation with the 561 nm laser (9.5 mW, 50%) through quad dichroic and emitted signal collected through emission filter EM-705/72. Laser powers measured after the objective.

**System B**

Nikon Ti-E inverted microscope equipped with a CFI Apo 60X/1.49NA oil-immersion objective lens and a 1.5X tube lens, a motorized stage (ProScan II, Prior Scientific), a scientific CMOS camera equipped with a 2048 x 2048 pixels (binned to 1024x1024 pixels) (Hamamatsu, Model #C15440) and a MLC400B (Keysight) laser including 561 and 640 nm lasers and filter cube containing 405/488/561/638 quad dichroic and barrier filters, an emission filter wheel with band pass filters detailed below (all filters, Chroma). Each image represents a 72 µm × 72 µm square region of the sample. The different channels can now be considered as a combination of incident laser wavelength and the corresponding bandpass filter. The “561 channel” consists of excitation with the 561 nm laser (9.4 mW, 70%) through quad dichroic and emitted signal collected through emission filter EM-603/50. The “640 channel” consists of excitation with the 640 nm laser (2.5 mW, 20%) and collected through quad dichroic and EM-705/72 emission filters. Laser powers measured after the objective.

**Calibration imaging experiments**

We calibrate each system regularly to determine channel offsets, illumination flatness, and regional point spread functions (PSF). For the channel offsets we dilute 100 nm Tetraspeck Fluorescent Microspheres (ThermoFisher, Cat #T7284), in 100uL methanol solvent and spotted them onto a glass slide to dry, adjusting the dilutions to achieve approximately 100 peaks per field. We captured images in all channels over 100 fields. Each microsphere contains dyes spanning multiple fluorescent channels and can be used to determine any fixed lateral offset between channel images. For the other metrics we use either the tetraspec calibration data or experimental samples of single count peptides.

We then analyze these images using the “calibration_sigproc” workflow, which, for multi-channel data, first performs a subpixel alignment via gradient descent after a fast Fourier transform to provide the per channel offsets. Next the peaks are identified in a similar manner as experimental data (detailed below in *signal processing*) however here each channel is treated independently. Using the calculated peak we parameterized the PSF, which models the shape and intensity of a single fluorescent peptide, by fitting each peak to 2d gaussians. The images were split into 25 sub regions and then the average PSF was calculated using all peaks and all fields, given a regional expectation for the PSF. Lastly, from the individual these fits we calculate the location and intensity of each peak which are used as a spatial indicator of illumination. Combining this information across all fields we arrive at a geometric expression of the regional illumination for each channel.

**Signal processing**

Signal processing includes the series of image processing steps converting multichannel images captured through the Nikon microscope (.nd2 files) after every Edman cycle into intensity arrays for each image channel across the cycles for every peptide spot. Glossary of terms used are shown in **Supplementary Table ST1**.

**Steps**

1. **Nd2 files are converted to npy files**. After every Edman cycle, we saved the images in Nikon's proprietary nd2 file, which comprises images from multiple channels and fields. Using an n2 converter python package, we converted the nd2 files (one per cycle, containing all channels for all fields) to numpy array files (one per field, containing all cycles for all channels). During this conversion process, we computed a per-channel/cycle field quality metric using a low-pass Fast Fourier Transform (FFT) filter to measure low-frequency power in the image. We later use this per-channel/cycle metric and average them to arrive at a field-quality measurement, which may be used for filtering ahead of classification.

   *Pseudo-code:*

   OUTPUT: numpy array (.npy file) per field, after reorganizing the contents of nd2 files, one per cycle.

   ```
   FOR every imaging cycle
       GET nd2 file that contains data for multiple channels and multiple fields
       OUTPUT. npy temporary files for each channel, cycle per field
       USING nd2 python library
   FOR every field
       GET all channel/cycle information from temporary. npy files for this field
       OUTPUT single. npy file containing all cycles/channels for this field
       USING nd2 python library
       COMPUTE and save low-frequency-power as a measure of field quality
   ```

2. **Regional illumination balancing is applied to account for variations in signal intensity over different regions within each image.** We balance each image to overcome non-uniform signal intensity (e.g. "vignetting" inherent when using spherical optics) based on the regional illumination measured during calibration experiments described above.

   *Pseudo-code:*

   OUTPUT: regionally balanced image as numpy array.

   ```
   FOR every field
       FOR every channel
           GET experimentally-determined balance_image for this channel
           FOR every cycle
               Divide channel-cycle image by balance_image
               SAVE regionally-balanced image
   ```

3. **Band-pass signal filtering**. We transform every regionally-balanced image into the frequency domain using a standard FFT algorithm (numpy) and reject signals above and below empirically-determined threshold frequencies to remove both background (reject-low) and over-saturated (reject-high) signals.

   *Pseudo-code:*
   ```
   OUTPUT: band-pass filtered image data
   FOR every field
       FOR every channel
           FOR every cycle
               USING regionally-balanced image
               REMOVE signal above and below reject thresholds via FFT filter
               SAVE filtered image
   ```

4. **Subpixel image-alignment, shift, and resample**. We align all images for a given field across cycles to account for stage movement between cycles. This involves, first an alignment done on one channel for all fields then the channel offsets, determined during calibration, is applied to the remaining channels. For the fixed channel alignment, we perform a first-pass pixel-level alignment via OpenCV's filter2d convolution, giving us pixel-offsets for each image relative to the first cycle's image. Then, we determine the sub-pixel offset using a gradient-descent in Fourier space to achieve sub-pixel accuracy. An alignment score for each field is calculated as the maximum shift in pixels required to align all system cycles, which we use downstream to filter out images prior to analysis.

   *Pseudo-code:*
   ```
   OUTPUT: resampled aligned images + alignment scores for every field stack
   FOR each field
       FOR each channel used for alignment
           FOR every cycle in the experiment
               ALIGN the filtered images in the field stack to the first one
               SAVE the pixels-shifted pair value per image (alignment score)
               USING pixels-shifted alignment offsets per image
               SHIFT image via FFT-based sub-pixel shifting
               RESAMPLE common region of interest from shifted images
   ```

5. **Find peaks *via* convolution.** We determine the locations of fluorescent peptides (peaks) for the first cycle because the signal must be present in at least the initial image to be a valid peptide signal. These peaks correspond to local maxima in signal intensity for each image. To find peaks with 1-pixel accuracy, we convolve an approximate point-spread-function kernel (an area under curve of 1.0 Gaussian that has been tuned to match observed empirical data) with the image in each channel. We then refine these locations to ½ pixel accuracy by using the center-of-mass of the already-identified peaks, determined by the regional context.

*Pseudo-code:*
OUTPUT: peak locations
FOR every channel:
FIND peaks in cycle 0 by convolving source image with approximate PSF
COMPUTE union of peaks across channels and return as a single list of locations

6. **Fit Gaussian parameters to some or all of the peaks.** We select at random a subset of the peaks found in step 5 and fit each peak to a 2D Gaussian. This serves to examine peak sizes, potentially at different cycles, as a proxy diagnostic for focus. Although this information is not used in the signal-processing pipeline "proper" (*i.e.* it is not an input to further downstream processing), it is displayed in reports viewed by persons analyzing the data by default.

7. **Compute radiometry parameters per peak.** We fit the parameters for the 2D Gaussian point-spread-functions during calibration to the peaks of control peptides located at various positions within the field. As discussed in calibration, the shape and intensity of these point-spread-functions depend on the peptide's location in the field. We use the appropriate parameters for the PSF based on a peak's location in the field to construct the expected PSF image ($PSF$), which acts as a kernel. This kernel is then convolved with the observed peak background removed image ($Data$) to derive the $signal$ (**Eq. S1b**) and $noise$ (**Eq. S1e**) for every peak in every channel and cycle.

$$\sigma_{PSF}^2 = \sum_i^{pixels} PSF_i^2 \qquad \text{(S1a)}$$

$$signal = \frac{\sum_i^{pixels} (PSF_i * Data_i)}{\sigma_{PSF}^2} \qquad \text{(S1b)}$$

$$e_i = Data_i - signal * PSF_i \qquad \text{(S1c)}$$

$$\sigma_e^2 = \sum_i e_i^2 \qquad \text{(S1d)}$$

$$noise = \sqrt{\frac{\sigma_e^2}{\sigma_{PSF}^2}} \qquad \text{(S1e)}$$

From the data we further calculate the number of cycles each peak remains fluorescent

*Pseudo-code:*
OUTPUT: peak information (radiometry)
FOR each peak
DEFINE kernel from PSF params based on location in field
CONVOLVE with peak
DETERMINE parameters such as signal and noise

8. **Collation of data into intensity reads and calculation of lifespan metrics.** We collate the peak information for the different channels and generate an intensity array for the channels associated with each peptide termed reads. From the data we are able to calculate the lifespan: the number of cycles each peak remains fluorescent. This is calculated from the minimum cosine distance between the measured reads and all possible unit normalized reads. It is the lifespan that is used to calculate the frequency histograms in **Figs. 2, S2, and S9**. We additionally calculate the intensity summary statistics for each peak during and after its lifetime.

9. **Information of all the identified peaks are collected for each channel across cycles and assembled as an intensity array associated with individual peptides.** We collate the peak information for the different channels and generate an intensity array for the channels associated with each peptide. We then combine these intensity reads into a multidimensional numpy array, which we call a radmat (radiometry matrix). In the radmat, every row is a peak and every column is the cycle, with signal and noise for the channels occupying different dimensions. We use a custom python dataframe to store information about radmat the other information about the peaks, and a separate data frame for all the metadata about the peak information including: field quality score, field alignment score, aligned position (x and y), and lifespan length. This flexible format allows for a number of computational transformations and extractions, and provides a list of all the information contained in the end of the data.

10. **Post Signal Processing Filtering:** For all post signal processing analyses, we remove poor quality reads using several filter metrics. First, we remove any fields where the alignment offset is greater than one third of a PSF sub region (150 pixels). This removes peaks having significant changes in illumination and/or PSF size from cycle to cycle. These extreme misalignments are rare with typical combined offsets between 5 and 25 pixels. Next we remove any field with poor field quality. As discussed above this value measures low frequency (large) structure in the image. Examples of these types of structures include large fluorescent contaminants (eg. dust, silane clusters, peptide aggregates) or large negative structures (e.g. bubbles). For consistency we set this value to 500 for all runs in this publication. To ensure that we are only analyzing single peptides we also filter by how well the peak resembles the expected PSF, i.e. low noise values. The most common cause of high noise is non-diffraction limited spots resulting from two or more peptides (or contamination) in close proximity. The noise threshold is chosen to reject above, approximately two standard deviations above mean noise distribution. Because noise and signal are correlated the noise threshold also increases with signal. Currently this threshold is set manually but we are working on methods for automation to improve reproducibility.

    Additional filtering may be used for specific analysis. For colocalization analysis a dark threshold is set at three sigmas above the background distribution. For FRET analysis and classification, low intensity contamination is removed by rejecting all peaks above the dark threshold and three sigmas below the mean one count intensity. Lastly for classification we also remove any high count anomalies at three sigmas above the highest count intensity distribution. We are working on removing both these high count and contamination anomalies from our workflow to avoid the need of this filtering in the future.

**Estimation of fluorosequencing parameters**

We divided the fluorosequencing parameters used in this publication into system-wide parameters and fluorophore-dependent parameters, which are defined in detail in our previous publications (Swaminathan, Boulgakov, and Marcotte 2015; Swaminathan et al. 2019). They are estimated here through a series of controlled experiments, parameter fitting, and estimations. Since the collection of this data presented here, we have developed automated methods of parameter estimation (Smith, 2023; Smith MB, VanderVelden K, Blom T, Stout HD, Mapes JH, Folsom TM, Martin C, Bardo AM, Marcotte EM. Estimating error rates for single molecule protein sequencing experiments. *bioRxiv* (2023) doi:10.1101/2023.07.18.549591) based on the whatprot classifier (Smith, Simpson, and Marcotte 2022) and other publicly available data fitting packages. The results in that publication show parameters consistent with the values presented here.

The system-wide parameters include the average probability of Edman failure (*p_edman*) and surface detachment rate (*p_detach*). Edman failure is the percent of molecules per cycle that do not undergo the removal of the N-terminal amino acid by Edman degradation. We modeled this in a similar method to that described in Swaminathan et al. 2019. As shown in this publication this value is highly dependent on both the experimental conditions and the peptide sequence and ranges from 1 to 20% per cycle. For the optimized conditions used in the classification experiments **(Fig. 4 and 5)**, a value of 5% per cycle was used in the classifier training simulations. During sequencing the entire peptides can be removed by either release of non-specifically bound peptides or hydrolysis of the underlying silane surface. The rate of this detachment from the surface (*p_detach*) is measured using peptides with two fluorophores and calculating the rate at which the signal for both fluorophores are lost in the same cycle. In our previous publication we reported values of 5% per cycle, here with the surface improvements we now measure this rate at 0.5% per cycle.

We determined the fluorophore-dependent parameters for each fluorophore, including Alexa555, TexasRed-X, and Atto643 (shown in **Supplementary Table ST2**). To determine these parameters, we conducted controlled experiments with dual-labeled peptides to calculate the surface detachment, above, and the dud-dye rate, and with N-terminally acetylated peptides (*JSP260, JSP229, JSP288*) which are not subject to Edman degradation chemistry, to isolate losses due to chemical-destruction.

To determine the per cycle photobleaching rate we continuously illuminate the peptide for 120 seconds and acquire an image every second. We plotted an exponential decay curve for the counts of single peptide molecules remaining over time. When acquiring the images, we used the imaging solvent (**Supplementary Table ST7)**. To determine the chemical destruction rate we imaged the acetylated peptides as with fluorosequencing after six Edman cycles and measured the loss rate of peaks per cycle. This measurement provides the combine per cycle loss due to photobleaching and chemical destruction rates (*p_bleach*).

As described previously, we observed that peptides with two counts of the same fluorophores, which was confirmed through mass characterization, appeared to have the brightness of only a single fluorophore. We speculate that the sample preparation process could have caused photobleaching or the formation/presence of a non-fluorescing isomer. To calculate the dud dye rate ($R_{dd}$, *p_dud*), we imaged multiple fields of the dual-labeled peptide. By measuring the count of peptides with one ($F_s$) and two ($F_d$) fluorophores, we computed the fraction of fluorophores that are considered "duds" using the **Equ. S2**. We note that this calculation will slightly underestimate the true dud dye rate as we are unable to calculate the fraction of peptides with no fluorescence.

$$R_{dd} = \frac{F_s}{2(F_s + F_d)} \qquad \text{(S2)}$$

Lastly, the mean intensity distribution and its standard deviation are calculated for the population of peptides with single fluorophore fluorescence.

**Building Machine Learning Classifier**

To infer peptide identity, we designed a workflow involving building a machine learning classifier which classifies and scores the signal data obtained from fluorosequencing directly to peptide identity.

1. **Reference peptide database is created**. We create an expected peptide database either from a protein list, simulating the peptides generated from protease digestion in the sample, or by generating directly from a list input peptide. In the case of building the four peptide classifier **Fig. 4**, we also include a random set of 50 peptides as a decoy list. In the case of the MHC peptide experiment, we use reference peptides identified using mass-spectrometry as the reference database. We then convert the peptide sequences to a “fluorostring” represented as [.0.1..1] where "." represents an unlabeled amino acid. The numbers represent the fluorophores for each channel (0 or 1).

2. **Using Monte Carlo simulations, synthetic fluorosequencing reads are generated for reference database peptides.** Using the experimentally obtained fluorosequencing parameters detailed above, we perform Monte-Carlo simulation for each peptide in the list by simulating 1000 copies for each peptide, labeling the selected amino acids with fluorophores, and using the probability of Edman failure, dud-dye, photobleaching, and dye-destruction, from the experimentally obtained fluorosequencing parameters detailed above, for the simulation. At each cycle, we simulate the possibility of Edman failure, dye bleaching, and so on, to arrive at a dye-sequence for the peptide. The resulting sequence is assigned a random value drawn from the intensity distribution for the channel dye, yielding the signal for the peptide at each cycle. Note that the information about the originating peptide is stored alongside the simulated fluorosequencing read. The sequence of radiometry at each cycle, in each channel, for each peptide follows the same format as the data produced by the instrument through the signal processing pipeline described above.

3. **A random forest classifier is trained on the synthetic reads.** We used the peptide/fluorosequencing data generated from Monte-Carlo simulation to construct a multi-class Random forest classifier. The number of features employed was determined by multiplying the number of channels and the number of cycles. Typically, the training set comprised 80% of the data, while the remaining 20% was reserved for testing.

**Raw intensity array data for each individual peptide obtained from fluorosequencing experiment is scored against the random forest Classifier.** We use the machine learning classifier to classify and score the intensity array (reads) generated from signal processing steps for each read. The classifier assigns a score to all peptide classes for each read, which can be considered a probability of assigning the read to the correct peptide class. The read is then attributed to the highest scoring peptide class. To determine a scoring threshold, we examine the scores associated with the decoy peptide list (known to be incorrect classifications) and the scores associated with the input peptides (known to be correct). In the case of the four peptide mixture samples (**Fig. 4**), we applied a score threshold of 0.99 and obtained the counts for the different classified peptides. In the case of the MHC peptide mixture samples (**Fig. 5**), we applied a score threshold of 0.7 and obtained the counts for the different classified peptides. For the high-scoring reads above, we also clustered the reads using the Python umap-learn package's default settings, indicating each read's assignment by color.
